# Corticostriatal suppression of appetitive Pavlovian conditioned responding

**DOI:** 10.1101/2021.08.19.456995

**Authors:** Franz R. Villaruel, Melissa Martins, Nadia Chaudhri

## Abstract

The capacity to suppress learned responses is essential for animals to adapt in dynamic environments. Extinction is a process by which animals learn to suppress conditioned responding when an expected outcome is omitted. The infralimbic cortex (IL) to nucleus accumbens shell (NAcS) neural circuit is implicated in suppressing conditioned responding after extinction, especially in the context of operant cocaine-seeking behaviour. However, the role of the IL-to-NAcS neural circuit in the extinction of responding to appetitive Pavlovian cues is unknown and the psychological mechanisms involved in response suppression following extinction are unclear. We trained rats to associate a 10 s auditory conditioned stimulus (CS; 14 trials per session) with a sucrose unconditioned stimulus (US; 0.2 mL per CS) in a specific context and then, following extinction in a different context, precipitated a renewal of CS responding by presenting the CS alone in the original Pavlovian conditioning context. Unilateral, optogenetic stimulation of the IL-to-NAcS circuit selectively during CS trials suppressed renewal. In a separate experiment, IL-to-NAcS stimulation suppressed CS responding regardless of prior extinction and impaired extinction retrieval. Finally, IL-to-NAcS stimulation during the CS did not suppress the acquisition of Pavlovian conditioning but was required for the subsequent expression of CS responding. These results are consistent with multiple studies showing that the IL-to-NAcS neural circuit is involved in the suppression of operant cocaine-seeking, extending these findings to appetitive Pavlovian cues. The suppression of appetitive Pavlovian responding following IL-to-NAcS circuit stimulation does not, however, appear to require an extinction-dependent process.

**SIGNIFICANCE STATEMENT:** Extinction is a form of inhibitory learning through which animals learn to suppress conditioned responding in the face of non-reinforcement. We investigated the role of infralimbic (IL) cortex inputs to the nucleus accumbens shell (NAcS) in the extinction of responding to appetitive Pavlovian cues and the psychological mechanisms involved in response suppression following extinction. Using *in vivo* optogenetics, we found that stimulating the IL-to-NAcS neural circuit suppressed context-induced renewal of conditioned responding after extinction. In a separate experiment, stimulating the IL-to-NAcS circuit suppressed conditioned responding in an extinction-independent manner. These findings can be leveraged by future research aimed at understanding how corticostriatal circuits contribute to behavioural flexibility and mental disorders that involve the suppression of learned behaviours.

## INTRODUCTION

The capacity to inhibit learned responses is essential for adaptive behaviour. Extinction is a fundamental psychological process by which animals learn to suppress responding to a conditioned stimulus (CS) that previously predicted a biologically significant unconditioned stimulus (US). New inhibitory learning is thought to occur during extinction when the CS is presented without the anticipated US (Konorski, 1948; Pearce and Hall, 1980). Following extinction, retrieval of this inhibitory memory suppresses conditioned responding. However, retrieval of the inhibitory extinction memory is context-dependent, and a renewal of conditioned responding can occur when the context is changed after extinction (Bouton, 1993; 2004). This impermanence of response suppression is a fundamental short-coming in using extinction to treat disorders such as post-traumatic stress and substance-abuse, which are characterized by heightened responses to environmental cues.

The infralimbic (IL) medial prefrontal cortex is a critical brain region for extinction (Quirk et al., 2000; Milad and Quirk, 2002; Rhodes and Killcross, 2004; 2007; Vidal-Gonzalez et al., 2006; Peters et al., 2008; 2009; Quirk and Mueller, 2008; LaLumiere et al., 2012; Do Monte et al., 2015; Villaruel et al., 2018; Lay et al., 2019) and is thought to mediate the extinction of appetitive and aversive conditioning through distinct projections to the nucleus accumbens shell (NAcS) and the basolateral amygdala (BLA), respectively (Peters et al., 2009; Bloodgood et al., 2018). Studies on the IL-to-NAcS neural circuit in appetitive extinction typically involve instrumental conditioning in which animals make an operant response to earn drug reinforcers. Pharmacologically disconnecting the IL and the NAcS reinstates extinguished cocaine-seeking (Peters et al., 2008), whereas promoting glutamatergic transmission between the IL and NAcS suppresses cocaine-seeking (LaLumiere et al., 2012). Furthermore, chemogenetic stimulation of the IL-to-NacS circuit reduces cue-induced cocaine-seeking (Augur et al., 2016). These results suggest that activity in the IL-to-NAcS circuit suppresses operant cocaine-seeking after extinction. Lastly, pharmacological disconnection of the IL and NAcS disrupts the capacity of a Pavlovian cue to invigorate operant responding, suggesting that the circuit is also involved in processing appetitive Pavlovian associations (Keistler et al., 2015). However, little is known about the role of the IL-to-NAcS neural circuit explicitly in extinction of appetitive Pavlovian conditioned responding.

The IL and NAcS may mediate the suppression of conditioned responding by facilitating the retrieval of an inhibitory extinction memory. Extinction increases IL activity (Milad and Quirk, 2002) and induces synaptic plasticity in the NAcS (Sutton et al., 2003). In aversive conditioning, pharmacologically stimulating the IL strengthens inhibitory memory and facilitates extinction retrieval (Lingawi et al., 2016; 2018). Furthermore, distinct neuronal ensembles that mediate the extinction of cocaine-seeking exist within the IL and IL projections to the NAcS (Warren et al., 2016; 2019). In some studies, extinction is a pre-requisite for IL and IL-to-NAcS circuit stimulation to suppress operant cocaine-seeking (Augur et al., 2016; Ewald Müller et al., 2019). However, optogenetic stimulation of the IL and the IL-to-NAcS circuit can also suppress operant food-seeking and cocaine-seeking without prior extinction training (Do Monte et al., 2015; Cameron et al., 2019). Therefore, it remains unclear if the IL-to-NAcS circuit suppresses responding through the retrieval and strengthening of an extinction memory, especially in appetitive conditioning.

Here, we examined the role of the IL-to-NAcS neural circuit in the suppression of appetitive Pavlovian conditioned responding after extinction. First, we tested whether optogenetic stimulation of the IL-to-NAcS circuit during presentations of a sucrose-predictive CS would suppress context-induced renewal of conditioned responding. Second, we adapted a procedure from Lingawi et al. (2016) to determine whether the suppression of conditioned responding to a sucrose-predictive CS was due to the retrieval of an inhibitory extinction memory that recruited the IL-to-NAcS neural circuit. Lastly, to determine if stimulating this circuit would non-specifically reduce behaviour, we investigated the effect of IL-to-NAcS stimulation on the acquisition and expression of appetitive Pavlovian conditioning. We predicted that stimulating the IL-to-NAcS circuit would suppress the return of appetitive Pavlovian conditioned responding by facilitating the retrieval of a previously established, inhibitory extinction memory.

## MATERIALS AND METHODS

### Subjects

Ninety-six male, Long-Evans rats (Charles River, Quebec, Canada: 220-240 g on arrival) were pair housed on arrival and single housed three days later. Rats were housed in polycarbonate home-cages (44.5 cm x 25.8 cm x 21.7 cm) containing Sani-Chip bedding (Envigo, 7090A), a nylon bone toy, (Bio-Serv, K3580) and a tunnel (Bio-Serv, K3245). Rats had unrestricted access to food (Charles River, 5075) and water in their home-cages throughout the experiment. Home-cages were in a colony room with controlled temperature (21°C) and humidity (44%) on a 12 h light/dark cycle (lights on at 7am). All procedures were conducted during the light phase. All procedures were approved by the Animal Research Ethics Committee of Concordia University and accorded with the guidelines from the Canadian Council on Animal Care.

### Apparatus

Behavioural procedures were conducted in six conditioning chambers (Med Associates, ENV-009A, St. Albans, VT, USA) housed in sound-attenuating, melamine cubicles. Chambers contained bar floors, a house light (75 W, 100 mA, ENV-215M) in the center of the left wall, and a white noise generator and speaker (Med Associates, ENV-225SM, 5 dB above background noise) in the top left corner of the left wall. A customized fluid port (ENV-200R3AM, opening height of 13.2 cm) was used to ease port access and was located 2 cm above the bar floor in the center of the right wall. Infrared sensors (ENV-254CB) flanked both sides of the port opening to detect port entries. Solutions were delivered into the fluid port via a polyethylene tube (Fisher Scientific, 141 691 A) connected to a 20 mL syringe in a pump (PHM-100, 3.33 rpm) located outside of the cubicle. All events, peripheral devices, and data collection were controlled by Med Associates software (Med-PC IV) on a computer.

Optogenetic stimulation was delivered in each conditioning chamber by a 150 mW, 473 nm laser (Shanghai Laser & Optics Century Co., BL473T3-150). The laser was connected to a unilateral optical rotary joint (Doric Lenses, FRJ-FC-FC, Quebec, Canada) via a 125 μm optical fiber (Fiber Optic Cable Shop, FC-FCFC-MS6-2M, Richmond, CA, USA). A custom-made patch cord (Trujillo-Pisanty et al., 2015) containing a 200 μm fiber connected the rotary joint to a custom-made optical fiber implant containing a 300 μm fiber. Prior to each test, the power of the laser was calibrated to approximately 30 mW for each optical fiber implant. Optogenetic stimulation was delivered at a frequency of 20 Hz (5 ms pulses) in a 10.2 s pulse train programmed through an Arduino microcontroller. Optogenetic parameters were based on our past study (Villaruel et al., 2018) and other published studies (Adamantidis et al., 2011; Britt et al., 2012).

### Solutions and Viruses

A 10% (w/v) sucrose solution was prepared by mixing sucrose (BioShop, 500070) in tap water and served as the unconditioned stimulus. Odours used in experiment 1 were prepared by diluting lemon oil (Lemon Odour; Sigma-Aldrich, W262528-1 KG-K) or benzaldehyde (Almond Odour; Fisher Scientific B240-500) with water to make a 10% solution. Viruses containing the transgene for channelrhodopsin 2 (ChR2) with the enhanced yellow fluorescent protein (eYFP) reporter (AAV5-CaMKIIa-hChR2(H134R)-EYFP, 1.5 x 10^13^ Vg/mL, Addgene) or eYFP alone (AAV2-CaMKIIa-EYFP, 2.0 × 10^12^ Vg/mL, Addgene; AAV5-CaMKIIa-EYFP, 9.0 × 10^12^ Vg/mL, Neurophotonics) were used for optogenetic surgeries.

### Surgery

Rats received stereotaxic surgery using standard procedures starting one week after single housing. Target coordinates for the IL were +2.9 mm anterior and +3.4 mm lateral relative to bregma and −5.8 mm ventral relative to the skull surface (30° angle). The viral vector containing the transgene for ChR2 or eYFP alone was microinfused into the IL (1 uL, 0.1 uL/min, 20 min diffusion). Microinfusion was conducted through a blunted 27 ¼ gauge needle (Fisher Scientific, 1 482 113B) connected via polyethylene tubing (PE20 VWR, CA-63 018-645) to a 10 uL Hamilton syringe (Hamilton, 1701N) on a pump (Pump 11 Elite, Harvard Apparatus, 704, 501). An optical fiber implant was inserted into the NAcS (AP +1.2 mm, ML +1.0 mm, from bregma and DV −7.5 mm from the skull surface) in the same hemisphere. Optical fiber implants were secured using five jeweller’s screws, Metabond (Patterson Dental, 5 533 484), and dental acrylic (A-M Systems; powder 525 000, solvent 526 000). Buprenorphine (Buprenex, 0.03mg/kg, s.c.) was administered as an analgesic after surgery. To facilitate recovery, rats were provided with banana-flavored oral rehydrator (PRANG, Bio-Serv, F2351-B) for 48 h post-surgery. Behavioural tests involving optogenetic stimulation were done at least 8 weeks after surgery to ensure ample time for virus expression.

Retrograde tracing with Cholera toxin subunit B (CTb; Invitrogen, ThermoFisher Scientific) was used to characterize the IL-to-NAcS and IL-to-BLA neural circuits. Rats (n=4) underwent stereotaxic surgery in which CTb conjugated with either Alexa Fluor 488 or Alexa Fluor 555 (0.5% weight/volume in 0.9% sterile saline) was unilaterally infused into the NAcS or BLA (0.3 uL, 0.1 uL/min, 10 min diffusion). Coordinates from bregma (AP and ML) and the skull surface (DV) for targeting the NAcS were AP +1.2 mm, ML +1.0 mm, DV −7.5 mm and for the BLA were AP −2.5 mm, ML −5.0 mm, DV −8.5 mm. The fluorescent label used for tracing was counterbalanced by region across rats.

### General Behavioural Procedures

Rats were handled and weighed before each procedure. Rats were exposed to 10% sucrose through a bottle in their home-cage in two 24 h sessions to acclimate them to the taste of sucrose. Rats were habituated to transport, the experimental room, and the conditioning chambers in a 20 min session in which rats were tethered to patch cords, the house light was on, and port entries were recorded. After this last habituation, rats received daily Pavlovian conditioning sessions (40 min). At 2 min after initiating the program, the house light was illuminated to signal the start of the session. Sessions consisted of 14 presentations of a 10 s continuous white noise CS occurring at a variable-time 140 s schedule (intertrial intervals 80, 140, or 200 s). Pumps were activated 4 s after CS onset to deliver 0.2 mL of sucrose into the fluid port (2.8 mL per session) and co-terminated with the CS. Ports were checked after each session to ensure sucrose consumption. Extinction sessions and tests occurred just as Pavlovian conditioning, but without sucrose. In all sessions, rats were tethered to a patch cord. However, patch cords were only functional during test sessions.

### Experiment 1: Effect of IL-to-NAcS stimulation on context-induced renewal of appetitive Pavlovian conditioned responding

Experiment 1 tested whether optogenetic stimulation of the IL-to-NAcS circuit during CS trials would attenuate context-induced renewal of appetitive Pavlovian conditioned responses after extinction. The conditioning context was referred to as Context A and the extinction context as Context B. The configuration of Contexts A and B were counterbalanced between two types. Context type 1 consisted of bar floors, black and white striped walls, and a lemon odour. Context type 2 consisted of wire grid floors, clear walls, and an almond odour. Odours were applied on petri dishes located underneath the floor of the conditioning chambers. Conditioning sessions occurred in Context A and extinction sessions occurred in Context B as previously described (Villaruel et al., 2018).

Groups consisted of rats microinfused with ChR2 (n = 10) or eYFP alone (n = 10). Rats received five successive habituation sessions to transport, the experimental room, being tethered to a patch cord in a default conditioning chamber, and the conditioning and extinction contexts. The order of habituation in the conditioning and extinction contexts was counterbalanced across rats. Habituation in the conditioning context involved fluid port training consisting of five un-signaled deliveries of sucrose with an inter-delivery interval of 240 s. After habituation, rats received 12 Pavlovian conditioning sessions in Context A, followed by at least three extinction sessions in Context B or until a criterion of 5 or fewer CS port entries was met. Following extinction, rats were tested in Contexts A and B across two days. Test order was counterbalanced such that half of the rats were tested in Context A first, and the other half in Context B first. Test sessions were separated by at least one extinction session or until the criterion of 5 or fewer CS port entries was met to mitigate any after-effects of optogenetic stimulation. We found that rats in the eYFP group that received their second test in Context A did not show renewal. Therefore, after test 2, all rats received two Pavlovian re-conditioning sessions in Context A and at least two extinctions in Context B or until the criterion was met prior to repeating the second renewal test. Final data analysis consists of collapsing the first renewal test and the repeated second test.

### Experiment 2: Effect of IL-to-NAcS stimulation on the suppression of appetitive Pavlovian conditioned responding after extinction

Experiment 2 tested whether prior extinction training was necessary for IL-to-NAcS stimulation to suppress appetitive Pavlovian conditioned responding. All behavioural sessions occurred in default conditioning chambers devoid of additional contextual cues. Rats received 10 daily sessions of Pavlovian conditioning. Next, rats microinfused with ChR2 (n = 25) or eYFP alone (n = 20) were divided into either an Extinction group (Ext) or No Extinction group (No Ext) matched on the acquisition of Pavlovian conditioning and CS port entries in the last conditioning session. Rats in the Extinction group (ChR2 n = 13; eYFP n = 10) received one extinction session 24 h after the last conditioning session. In contrast, rats in the No Extinction group (ChR2 n = 12; eYFP n = 10) did not receive extinction training and were instead handled and weighed in the colony room. The following day, all rats underwent a reconditioning session to re-establish baseline responding. An extinction test (Test 1) was conducted the following day and was identical to an extinction session but with optogenetic stimulation delivered during CS trials. An extinction retrieval test (Test 2) was conducted the next day and was identical to an extinction session but occurred without the delivery of optogenetic stimulation.

### Experiment 3: Effect of IL-to-NAcS circuit stimulation on the acquisition and expression of appetitive Pavlovian conditioning

Experiment 3 tested whether IL-to-NAcS stimulation would indiscriminately suppress CS responding, thereby preventing the acquisition of appetitive Pavlovian conditioning. All behavioural sessions occurred in default conditioning chambers. Rats (ChR2 n = 11, eYFP n = 11) received 12 daily sessions of Pavlovian conditioning as previously described with the exception that optogenetic stimulation was delivered during CS trials.

Following conditioning, we examined the effect of IL-to-NAcS stimulation on the expression of conditioned responding. Rats were tested across two sessions approximately 24 h apart for the expression of conditioned responding to the CS alone in the absence of the sucrose US. In one test optogenetic stimulation during the CS was present and in the other test optogenetic stimulation was withheld. Test order was counterbalanced across rats and rats were matched based on acquisition of Pavlovian conditioning measured by Δ CS port entries.

### Histology

All rats were euthanized with a pentobarbital (Euthanyl™) overdose and transcardially perfused with 0.1M phosphate buffered saline (PBS) followed by 4% paraformaldehyde (PFA) in 0.1M phosphate buffer. Brains were extracted and post-fixed in 4% PFA solution for 24 h followed by a 30% sucrose solution for 48 h in 4 °C. Brains were frozen at − 80 °C and sectioned using a cryostat (40 μm) in a one-in-five series. Brain sections were mounted onto microscope slides and processed for Nissl staining or fluorescence microscopy with DAPI (Vector labs, H-1200) to verify optical fiber placement and transgene expression. Images of transgene expression were captured through an epifluorescence microscope (Nikon Eclipse TiE) using a 4x lens for cell bodies and a 20x lens for neuron terminals. A rat brain atlas (Paxinos and Watson, 2007) was used to approximate the location of sections relative to bregma and the images were used to model the spread of transgene expression in the IL (Adobe Illustrator).

Rats in the retrograde tracing experiment were euthanized one week after receiving surgery and brains were processed as described above. Brain sections were stained with DAPI, cover slipped, and processed through fluorescence microscopy. Infusion sites were examined to ensure accurate targeting of the NAcS and the BLA using an epifluorescence microscope with a 4x lens. A confocal laser scanning microscope (Nikon C2) was used to image CTb labelled cells in the medial prefrontal cortex (4 sections per rat) using a 20x lens. Images were imported to Imaris Cell Imaging Software (Bitplane, Oxford Instruments) in which analysis was specifically restricted to the IL by the experimenter using a rat brain atlas (Paxinos and Watson, 2007). CTb-488 and CTb-555 labelled cells were defined using the ‘Blobs’ tool in Imaris, local contrast thresholding, and volume of labelled pixels. Co-labelling of CTb-488 and CTb-555 signals was determined as objects labelled in one channel that had greater than 20% of their volume also labelled in the second channel. The number of labelled and co-labelled cells was averaged across the 4 sections to get a single value for each rat. Density of labelled and co-labelled cells was calculated by dividing the average number of labelled and co-labelled by the average area of the selected quantified region across 4 brain sections.

### Fos Immunohistochemistry

Rats in Experiment 1 received an additional test to induce c-Fos and verify that optogenetic stimulation of IL-to-NAcS terminals expressing ChR2 had a physiological effect (Fuchikami et al., 2015; Benn et al., 2016; Wood et al., 2019). The c-Fos induction session occurred in default conditioning chambers and was identical to previous test sessions. Optogenetic stimulation was delivered for 14 trials to mimic previous tests, but in the absence of house light illumination, the white noise CS, or sucrose. Rats remained in the conditioning chambers for an additional 50 min to ensure that they were euthanized 90 min after the start of the session to maximize c-Fos expression (Muller et al., 1984; Bossert et al., 2011; Warren et al., 2016). Brain sections were processed in an anti-cFos rabbit antibody (1:2000; Cell Signalling, 2240) for approximately 72 h, and subsequently in a secondary solution with biotinylated goat anti-rabbit antibody (1:250; Vector Labs, BA-1000). Next, sections were placed in a tertiary of avidin and biotinylated horseradish peroxidase (1:1000; ABC kit, Vector Labs, PK-6100) and stained with a 3, 3’-diaminobenzidine (DAB) solution. Finally, sections were rinsed in phosphate buffer, mounted on slides, and cover slipped. Images of each section were captured through a brightfield microscope (Nikon Eclipse TiE) using a 10x lens. Two sections from the IL and the NAcS were chosen for quantification based on location relative to bregma and image quality. A rat brain atlas (Paxinos and Watson, 2007) was used to approximate the location of sections relative to bregma and the regions of interest. Image analysis was done through ImageJ FIJI. A region of the IL and the NAcS was selected manually for each section in both the stimulated and non-stimulated hemisphere. Quantification of the selection was done through a custom-made FIJI macro, which counted Fos positive nuclei based on colour relative to background, size, and circularity. Counts were then divided by the average area selected in FIJI to calculate density. The final Fos density for each rat consisted of the average across two sections for each hemisphere and region.

### Statistical Tests

Data was processed in Microsoft Excel, visualized in Prism (Graphpad), and analyzed in SPSS (IBM). Behavioural data of interest included port entries made during the entire session (total), 10 s prior to CS onset (Pre CS), during the 10 s CS (CS), 10 s after CS onset (Post CS), and during the intertrial intervals (between post CS offset and pre CS onset). Conditioned responding was measured using a difference score (Δ CS Port Entries) calculated by subtracting port entries made during the Pre CS period from the CS period to account for variability in baseline activity (Rhodes and Killcross, 2004; 2007; Villaruel et al., 2018). CS port entries were also analyzed per block consisting of two CS trials or per CS trial. Probability, duration, and latency of CS port entries were also collected and analyzed both as an average in the session and per CS trial. Probability was calculated as the number of trials with a port entry divided by the total number of trials (14). Duration was measured as time in the port after initiating a port entry during the CS. Latency was measured as time to initiate the first CS port entry. All significant interactions were further examined with Bonferroni corrected comparisons. The alpha level for statistical significance was set to p < 0.05.

## RESULTS

### Histology

We used a retrograde tracer (Cholera toxin B) to characterize neural projections from the IL-to-NAcS (Figure 1A-D). In the same rats we also examined the IL-to-BLA circuit, which is implicated in extinction of aversive Pavlovian conditioning (Peters et al., 2009; Bloodgood et al., 2008). Neural tracing of IL projections to the NAcS and BLA revealed largely distinct, non-overlapping projections to these output regions. Only a small proportion of labelled cells were found to project to both the NAcS and the BLA (Figure 1D).

**Figure 1.**
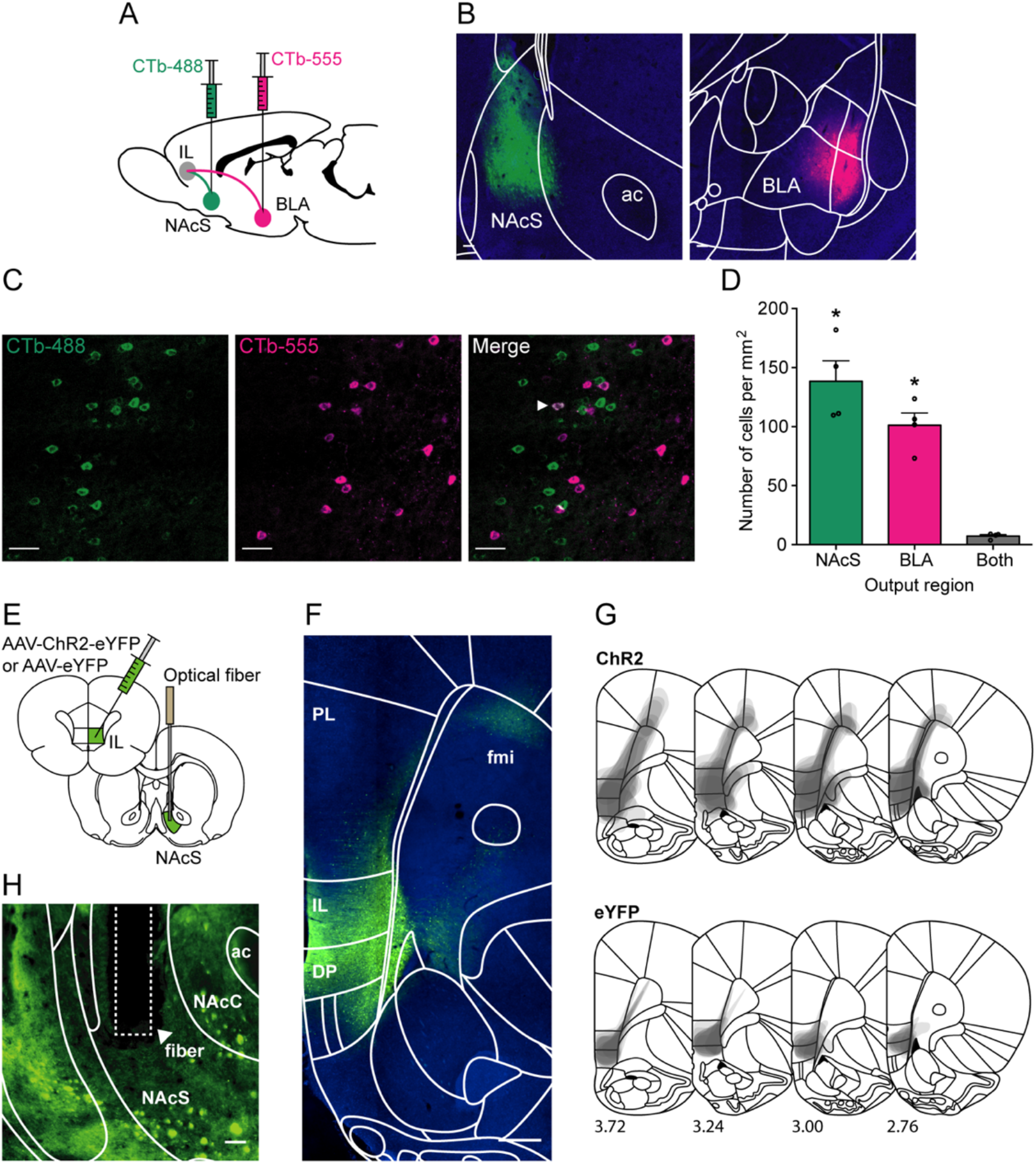
Neural tracing and optogenetic targeting of the IL-to-NAcS circuit. (**A)** Method schematic for neural tracing. CTb-488 and CTb-555 retrograde tracers were injected in the NAcS and the BLA, and quantification of labelled cells was done in the IL. (**B**) Representative images of injection sites in the NAcS (left) and the BLA (right). Anterior commissure (ac). Scale bar 100 μm. (**C**) Representative images of labelled cells in the IL. Arrow in merged image shows an example of co-labelling. Scale bar 50 μm. (**D**) Quantification of labelled cells in the IL shows largely non-overlapping cells projecting to the NAcS and the BLA. Data presented as mean ± SEM. * p < 0.05 vs. Both output regions. (**E)** Method schematic of microinjections of ChR2 or eYFP alone in the IL and an optical fiber implanted in the NAcS. **(F)** Representative image of ChR2 expression in the IL. Prelimbic cortex (PL), dorsal peduncular cortex (DP), forceps minor of the corpus callosum (fmi). Scale bar 500 μm. **(G)** Schematic depicting the extent of ChR2 (n = 9, top panel) and eYFP alone (n = 9, bottom panel) expression in the IL across four bregma points. **(H)** Representative image depicting ChR2 expression in IL terminals and the optical fiber within the NAcS. Nucleus accumbens core (NAcC), anterior commissure (ac). Scale bar 100 μm.

Figure 1E depicts the method for targeting the IL-to-NAcS neural circuit using optogenetics. Expression of the ChR2 transgene was observed in the IL (Figure 1F) and in terminals in the NAcS (Figure 1H). The approximate spread of the transgenes for ChR2 and eYFP alone was based on rats in Experiment 1 but was consistent across experiments (Figure 1G). The highest concentration of ChR2 expression was in the infralimbic cortex, the dorsal peduncular cortex and the ventral regions of the prelimbic cortex. Some expression of ChR2 was observed along the injector tract, in the anterior and lateral areas of the prelimbic cortex along the forceps minor of the corpus callosum and the anterior medial and ventral orbitofrontal cortex. A representative image of correct optical fiber placement in the NAcS is shown in Figure 1H.

In all experiments, rats were excluded from final behavioural data analysis due to lack of transgene expression or misplacement of optical fiber implants. In experiment 1, two rats (eYFP n = 1, ChR2 n = 1) were excluded due to misplaced optical fiber implants. The final group sizes for experiment 1 were n = 9 ChR2 and n = 9 eYFP. One additional rat (eYFP n = 1) from experiment 1 was removed from c-Fos immunohistochemistry analysis due to complications with histology. In Experiment 2, two rats (eYFP n = 1, ChR2 n = 1) were excluded due to misplaced optical fiber implants and one rat (eYFP n = 1) was excluded due to lack of transgene expression. The final group sizes for experiment 2 were n = 13 ChR2-Ext, n = 12 ChR2-No Ext, n = 10 eYFP-Ext, and n = 10 eYFP-No Ext. The final group sizes for experiment 3 were n = 11 ChR2, n = 11 eYFP.

### IL-to-NAcS stimulation induced Fos reactivity in the IL and NAcS

Fos immunohistochemistry was conducted on a subset of rats from experiment 1 (ChR2 n = 9, eYFP n = 8) to verify that optogenetic stimulation of the IL-to-NAcS circuit activated the IL and the NAcS (Figure 2A). In the IL (Figure 2B, left panel), Fos immunoreactivity was greater in rats expressing ChR2 than eYFP alone (Figure 2C, left panel; Virus, F(1,15) = 40.70, p < .001) and in the stimulated hemisphere relative to the non-infected, non-stimulated, control hemisphere (Hemisphere, F(1,15) = 9.50, p = .008). Density of Fos positive nuclei in the IL showed a statistically significant interaction between virus and hemisphere (Hemisphere x Virus, F(1,15) = 9.68, p = .007). The stimulated hemisphere had greater Fos immunoreactivity than the non-stimulated hemisphere in the ChR2 group (p < .001) but not in the eYFP group (p = .984).

**Figure 2.**
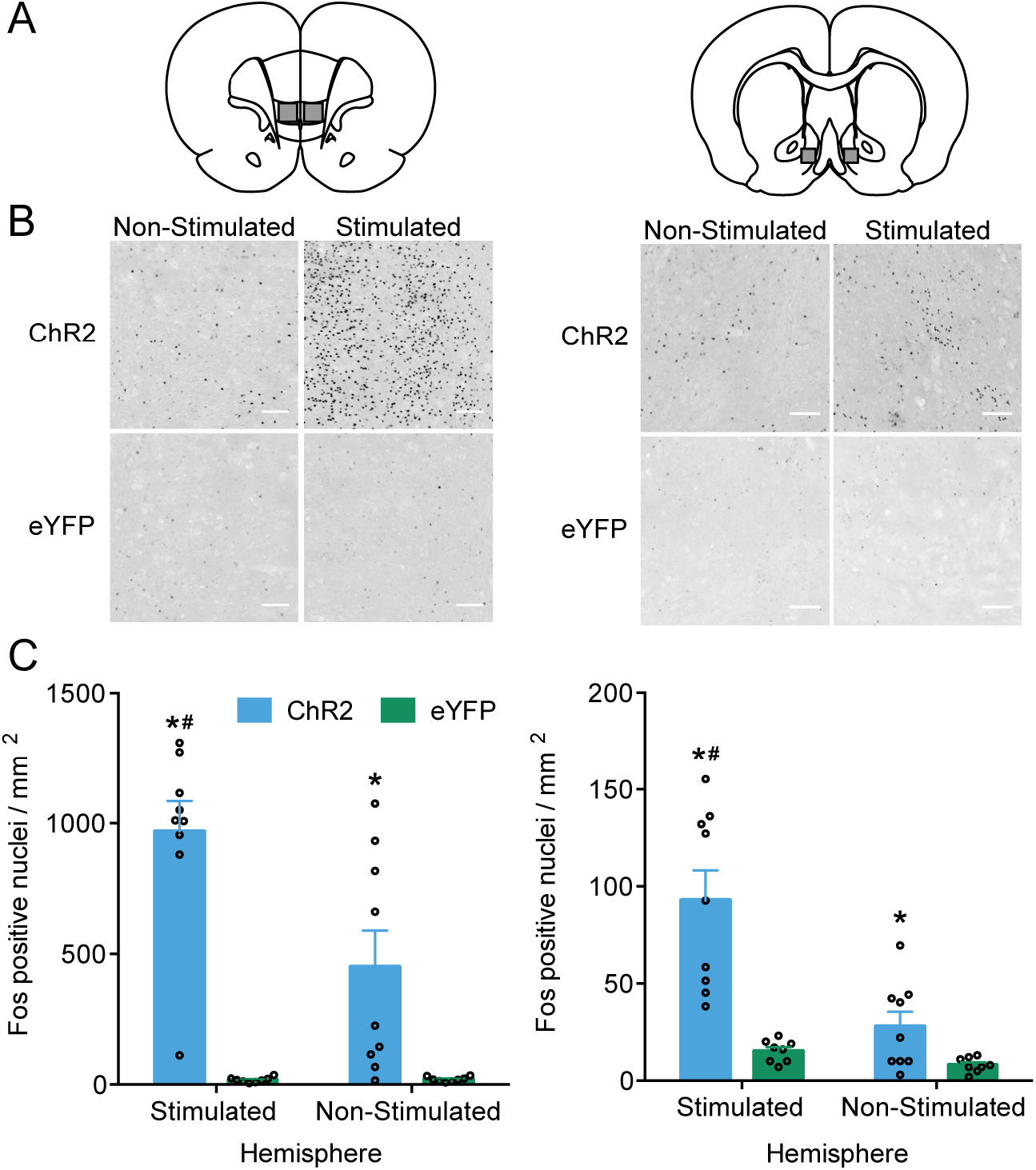
Quantification of Fos positive nuclei density following optogenetic stimulation of the IL-to-NAcS circuit. **(A)** Schematic depicting demarcations in the images of the IL (left panel) and NAcS (right panel) quantified for Fos positive nuclei. **(B)** Representative images of Fos positive nuclei in the IL (left panel) and NAcS (right panel) in the stimulated and non-stimulated hemisphere of rats expressing ChR2 or eYFP alone. Scale bars 100 μm. **(C)** Density of Fos positive nuclei (mean ± SEM) in the IL (left graph) and the NAcS (right graph) in rats expressing ChR2 or eYFP alone in both the stimulated hemisphere containing the optical fiber and in the non-stimulated hemisphere without an optical fiber. * p < 0.05 ChR2 vs. eYFP in each hemisphere. # p < 0.05 stimulated vs. non-stimulated hemisphere in the ChR2 group. All data are mean ± SEM.

However, the ChR2 group had greater Fos density than the eYFP group in both the stimulated hemisphere (p < .001) and the non-stimulated hemisphere (p = .011). These results indicate that optogenetic stimulation of IL neuron terminals in the NAcS activated the IL. However, optogenetic stimulation of the ChR2-transfected hemisphere also activated the opposite, non-stimulated hemisphere.

In the NAcS (Figure 2B, C, right panel), Fos immunoreactivity was greater in rats expressing ChR2 than eYFP alone (Virus, F(1,15) = 20.26, p < .001) and in the stimulated hemisphere relative to the non-stimulated hemisphere (Hemisphere, F(1,15) = 27.39, p < .001). A statistically significant interaction was observed in Fos density in the NAcS (Hemisphere x Virus, F(1,15) = 17.60, p = .001). Greater Fos immunoreactivity was observed in the stimulated hemisphere relative to the non-stimulated hemisphere in the ChR2 group (p < .001), but not in the eYFP group (p = .486). Fos density in the ChR2 group was also greater than the eYFP group in both the stimulated (p < .001) and non-stimulated hemisphere (p = .026). Therefore, optogenetic stimulation of IL neuron terminals in the NAcS activated the NAcS. As with the IL, optogenetic stimulation of the ChR2-transfected hemisphere also induced moderate activation in the non-stimulated hemisphere in the NAcS.

### IL-to-NAcS stimulation suppressed context-induced renewal of appetitive Pavlovian conditioned responding

Experiment 1 tested whether optogenetic stimulation of the IL-to-NAcS circuit would suppress the renewal of appetitive Pavlovian conditioned responding after extinction. Figure 3A depicts the optical fiber placement in the NAcS for all rats included in behavioural data analysis. Rats received Pavlovian conditioning in Context A, extinction in Context B, followed by two counterbalanced tests in Contexts A and B (Figure 3B).

**Figure 3.**
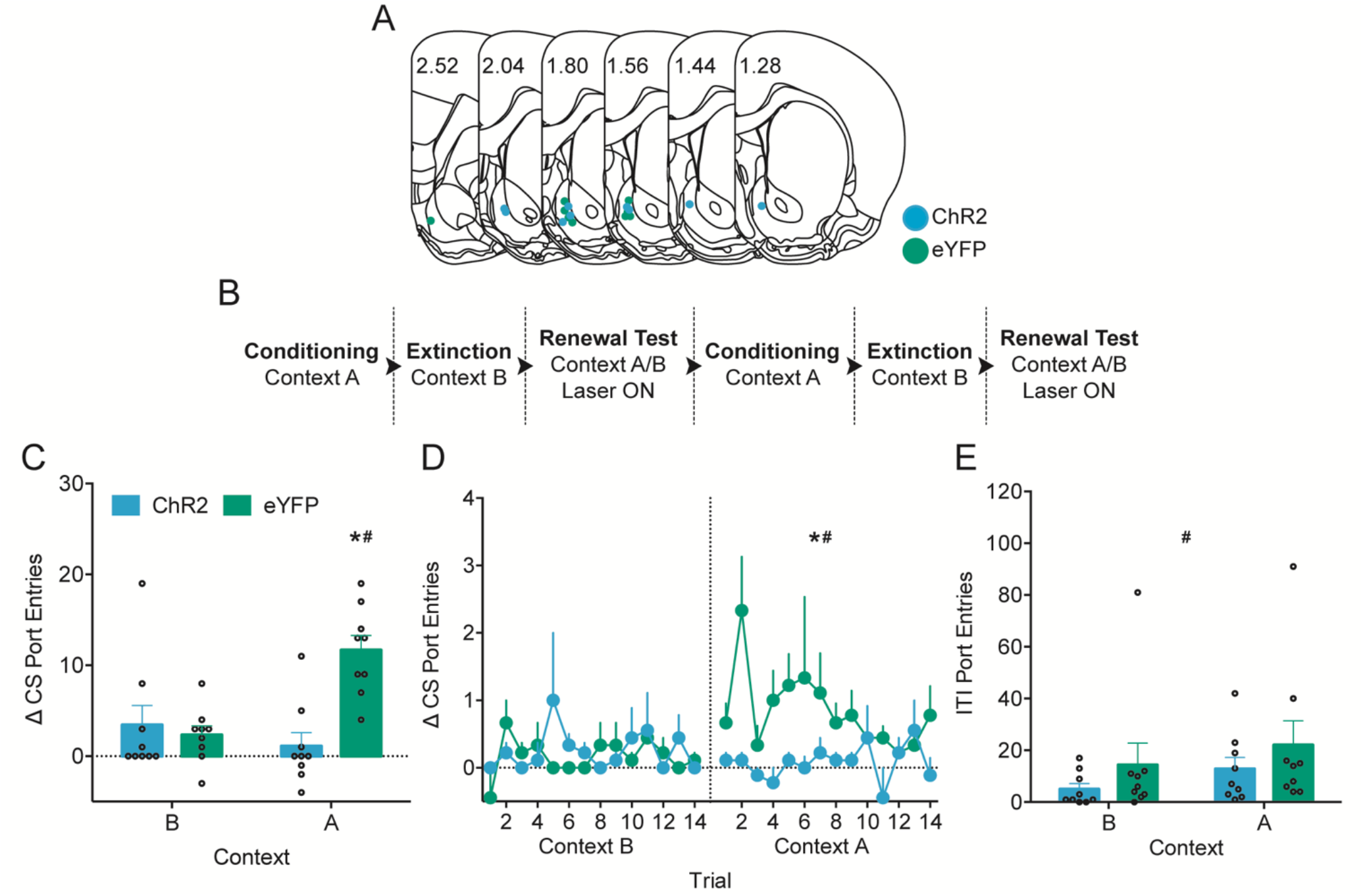
Optogenetic stimulation of the IL-to-NAcS neural circuit suppressed renewal of appetitive Pavlovian conditioned responding. **(A)** Optical fiber placements in the NAcS for ChR2 (blue) or eYFP alone (green) expressing rats included in the final data analysis. Numbers are locations of sections relative to bregma. **(B)** Design of behavioural procedures. **(C)** Δ CS port entries at tests in the conditioning context (Context A) for renewal relative to the extinction context (Context B). # p < 0.05 Context A vs. Context B in the eYFP group. * p < 0.05 ChR2 vs eYFP in Context A. **(D)** Δ CS port entries across trials at tests in Context A for renewal and the extinction context, Context B. # p < 0.05 Context A vs. Context B in the eYFP group across trials. * p < 0.05 ChR2 vs. eYFP in Context A across trials. **(E)** ITI port entries during tests in Context A and Context B. # p < 0.05 main effect of context. All data are mean ± SEM.

Optogenetic stimulation of the IL-to-NAcS during CS trials in the ChR2 group suppressed context-induced renewal of appetitive Pavlovian conditioned responding (Figure 3C). In the renewal tests, the eYFP but not the ChR2 group showed a robust renewal of conditioned responding in Context A relative to Context B (Context, F(1,16) = 18.38, p = .001; Virus, F(1,16) = 4.96, p = .041; Virus x Context, F(1,16) = 51.04, p < .001). Δ CS port entries were low for both eYFP and ChR2 at test in the extinction Context B (p = .644). However, Δ CS port entries were greater in the eYFP group compared to the ChR2 group in Context A (p < .001) indicating that stimulation of the IL-to-NAcS during CS trials suppressed renewal. The eYFP group showed greater Δ CS port entries at test in Context A relative to Context B (p < .001). In contrast, the ChR2 group showed similar levels of responding in both Context A and B (p = .060). A similar pattern of results was observed in measures of probability, duration, and latency of CS port entries (Supplementary Figure 1).

Analysis of Δ CS port entries across trials at test showed that IL-to-NAcS stimulation during CS trials suppressed conditioned responding in all trials (Figure 3D; Context, F(1,16) = 18.38, p = .001; Virus: F(1,16) = 4.96, p = .041; Context x Virus, F(1,16) = 51.04, p < .001; Trial, F(13,208) = 1.37, p = .240; Trial x Virus, F(13,208) = 1.49, p = .200; Context x Trial, F(13,208) = .68, p = .620.; Context x Trial x Virus: F(13,208) = .83, p = .519). Δ CS port entries were similar for ChR2 and eYFP groups in Context B (p = .644). However, at test in Context A, the eYFP group had greater CS port entries relative to the ChR2 group (p < .001). The eYFP group exhibited renewal and showed greater Δ CS responding in Context A relative to Context B across trials (p < .001). In contrast, the ChR2 group had equivalent responding across all trials in Contexts A and B (p = .060).

These results suggest that optogenetic stimulation of the IL-to-NAcS circuit suppressed appetitive Pavlovian conditioned responding throughout the renewal test. Altogether, optogenetic stimulation of the IL-to-NAcS neural circuit during CS trials attenuated the renewal of appetitive Pavlovian conditioned responding.

Stimulation of the IL-to-NAcS neural circuit did not affect port entries made during the intertrial intervals (ITI, Figure 2E). Port entries made during the ITI were greater in Context A than Context B in both ChR2 and eYFP groups (Context, F(1,16) = 6.24, p = .024; Virus, F(1,16) = 1.05, p = .322; Virus x Context, F(1,16) < 0.01, p = .986). Therefore, optogenetic stimulation of the IL-to-NAcS circuit did not produce non-specific motor effects during time intervals outside the CS and stimulation.

In a separate set of rats (n=6) we found that optogenetic stimulation of the IL-to-BLA neural circuit had no effect on the renewal of appetitive Pavlovian conditioned responding (Supplementary Figure 2 and 3).

### IL-to-NAcS stimulation suppressed appetitive Pavlovian conditioned responding regardless of prior extinction

Experiment 2 tested whether prior extinction training and the establishment of an inhibitory extinction memory were necessary for optogenetic stimulation of the IL-to-NAcS to suppress appetitive Pavlovian conditioned responding. Figure 4A depicts the optical fiber placement in the NAcS for rats included in the behavioural data analysis. The timeline of behavioural procedures is depicted in Figure 4B. All rats received Pavlovian conditioning, followed by either a single extinction session or handling, and then received a single reconditioning session to re-establish baseline responding (Supplementary Figure 4). Following reconditioning, all rats were tested for expression of appetitive Pavlovian conditioned responding under extinction conditions (Test 1).

**Figure 4.**
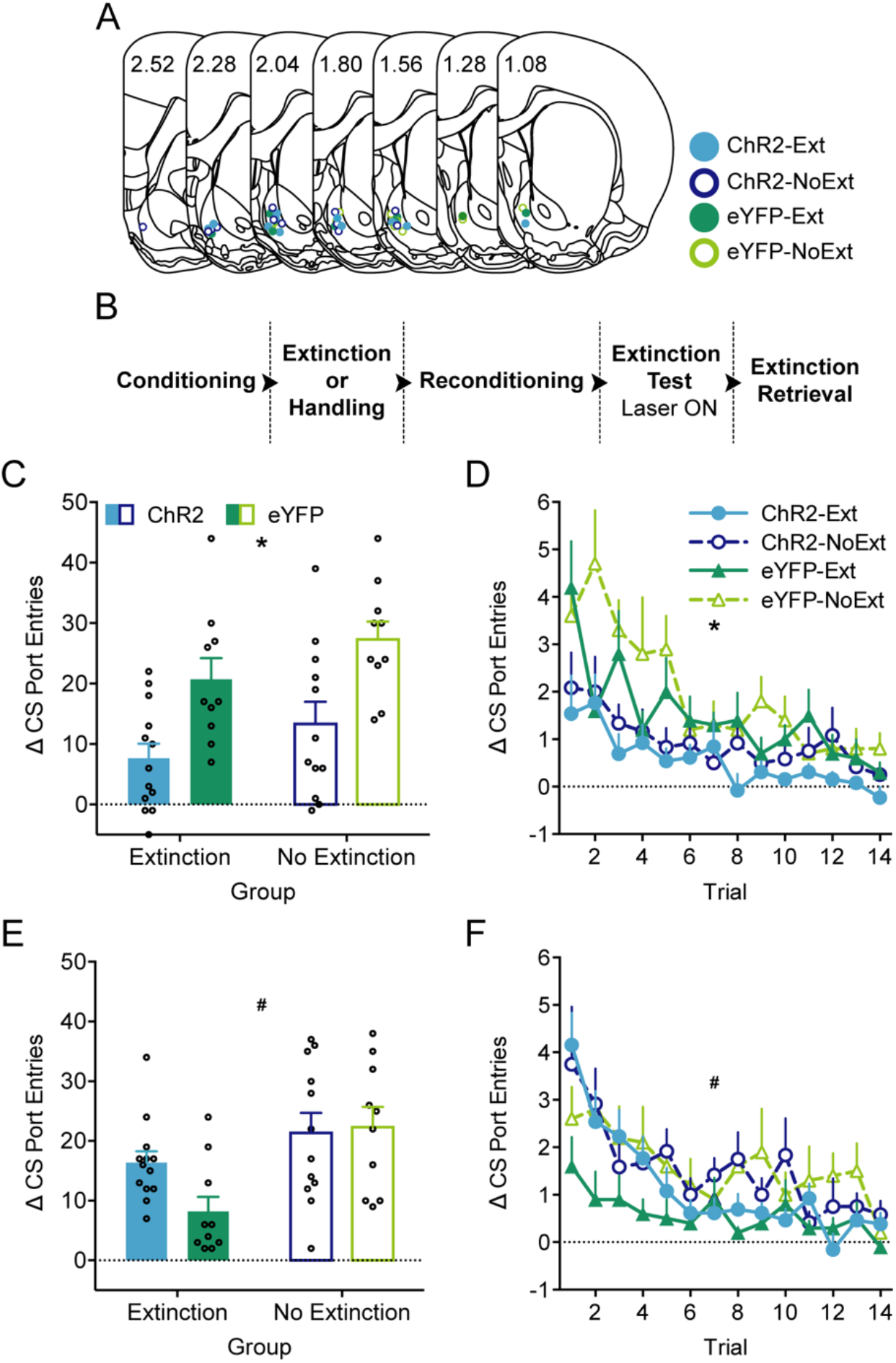
Optogenetic stimulation of the IL-to-NAcS neural circuit suppressed Δ CS port entries regardless of prior extinction and did not facilitate extinction retrieval. **(A)** Optical fiber placements in the NAcS of rats expressing either ChR2 (blue) or eYFP alone (green) in the extinction (filled) or no extinction (open) group included in the final data analysis. Numbers are locations of sections relative to bregma. **(B)** Design of behavioural procedures. **(C)** Δ CS port entries during the extinction test (Test 1) with IL-to-NAcS stimulation during CS trials. **(D)** Δ CS port entries across trials during the extinction test (Test 1) with IL-to-NAcS stimulation during the CS. **(E)** Δ CS port entries during the extinction retrieval test (Test 2) without IL-to-NAcS stimulation. **(F)** Δ CS port entries across trials during the extinction retrieval test (Test 2) without IL-to-NAcS stimulation. **(C-E)** * p < 0.05 main effect of virus group. # p < 0.05 main effect of extinction group. All data are mean ± SEM.

In Test 1, optogenetic stimulation of the IL-to-NAcS neural circuit during CS trials suppressed Δ CS port entries in the ChR2 group relative to the eYFP group regardless of prior extinction training (Figure 4C; Virus, F(1,41) = 18.32, p < .001; Group, F(1, 41) = 3.80, p = .058; Virus x Group, F(1, 41) = 0.02, p = .890). A similar pattern of results was observed in probability, duration, and latency of CS port entries, while ITI port entries were not affected by optogenetic stimulation during the CS (Supplementary Figure 5).

Δ CS port entries decreased across trials within the test, but were overall greater in the eYFP group relative to the ChR2 group (Figure 4D; Trial, F(13,533) = 10.71, p < .001; Virus, F(1,41) = 18.32, p < .001; Group, F(1,41) = 3.80, p = .058; Virus x Group, F(1,41) = 0.02, p = .890). There was no statistically significant interaction between trial, virus, and group (Trial x Virus, F(13,533) = 1.67, p = .129; Trial x Group, F(13,533) = 1.04, p = .400, Trial x Virus x Group, F(13,533) = 1.45, p = .197).

Therefore, stimulation of the IL-to-NAcS circuit suppressed appetitive Pavlovian conditioned responding regardless of prior extinction training and did so from the very first trial.

We conducted another extinction session (Test 2) the following day in the absence of optogenetic stimulation to determine if prior stimulation of the IL-to-NAcS neural circuit would facilitate extinction retrieval of appetitive Pavlovian conditioned responding. This prediction was based on findings in aversive Pavlovian conditioning studies in which stimulation of the IL during extinction facilitated subsequent extinction retrieval (Milad and Quirk, 2002; Milad et al., 2004; Do Monte et al., 2015; Lingawi et al., 2016; 2018).

In Test 2, Δ CS port entries were lower in rats that received prior extinction training relative to the No Extinction group (Figure 4E; Group, F(1, 41) = 8.71, p = .005) with no differences between ChR2 and eYFP groups (Virus, F(1, 41) = 1.75, p = .193). There was a near significant interaction (Virus x Group; F(1, 41) = 3.24, p = .079), supporting a possible effect of prior IL-to-NAcS stimulation on extinction retrieval. Exploratory simple effect comparisons showed a significant difference between ChR2 and eYFP groups within the Extinction group (F(1,41) = 4.97, p = .031) but not in the No Extinction group (F(1,41) = .11, p = .740). Visual inspection of Figure 4E shows that within the Extinction group, Δ CS port entries were higher in the ChR2 group than in the eYFP group, suggesting impaired extinction retrieval in the ChR2 Extinction group. Furthermore, Δ CS port entries were lowest in the eYFP group that had previously received extinction compared to all other groups. Similarly, within the Extinction group, the ChR2 group had a higher probability and duration of CS port entries than the eYFP group (Supplementary Figure 6). Furthermore, within the Extinction group, latency of CS port entries was statistically greater in the ChR2 group compared to the eYFP group (Supplementary Figure 6). ITI port entries were unaffected and similar across all four groups at test (Supplementary Figure 6).

Δ CS port entries decreased across trials (Figure 4F; Trial, F(13,533) = 9.06, p < .001) but were greater in the No Extinction group relative to the Extinction group during the extinction retrieval test (Group, F(1,41) = 11.33, p = .002, Virus, F(1,41) = 1.63, p = .209, Virus x Group, F(1,41) = 2.61, p = .114). This effect was largely mediated by the difference in the eYFP Extinction and eYFP No Extinction groups, as both ChR2 groups performed similarly across trials. There were no statistically significant interactions between trial, virus, and group (Trial x Virus, F(13,533) = 1.55, p = .147; Trial x Group, F(13,533) = .50, p = .844, Trial x Virus x Group, F(13,533) = 0.92, p = .498). Exploratory analysis comparing ChR2 and eYFP Extinction groups alone revealed that the ChR2 group made significantly more Δ CS port entries than the eYFP group at Test 2 (Virus, F(1,21) = 7.19, p = .014). Δ CS port entries decreased across Trial (F(13,273) = 7.22, p < .001), with a significant Trial x Virus interaction (F(13,273) = 2.31, p = .041) Additional post-hoc analysis indicate that within the Extinction group, Δ CS port entries were higher in the ChR2 group than in the eYFP group in Trial 1 (ChR2 vs eYFP, p = .013).

Together, these results suggest that prior IL-to-NAcS stimulation did not facilitate extinction retrieval, and exploratory analyses suggest that stimulation may have instead impaired extinction retrieval in rats that previously received extinction training.

### IL-to-NAcS circuit stimulation did not prevent the acquisition of Pavlovian conditioned responding

Experiment 3 tested whether IL-to-NAcS stimulation during CS trials would lead to general response inhibition and prevent the acquisition of appetitive Pavlovian conditioning. Figure 5A depicts the optical fiber placement in the NAcS for rats included in the behavioural data analysis. The timeline of behavioural procedures is depicted in Figure 5B. Rats received Pavlovian conditioning with optogenetic stimulation of the IL-to-NAcS circuit during CS trials followed by two counterbalanced expression tests under extinction conditions with either the presence or absence of stimulation.

**Figure 5.**
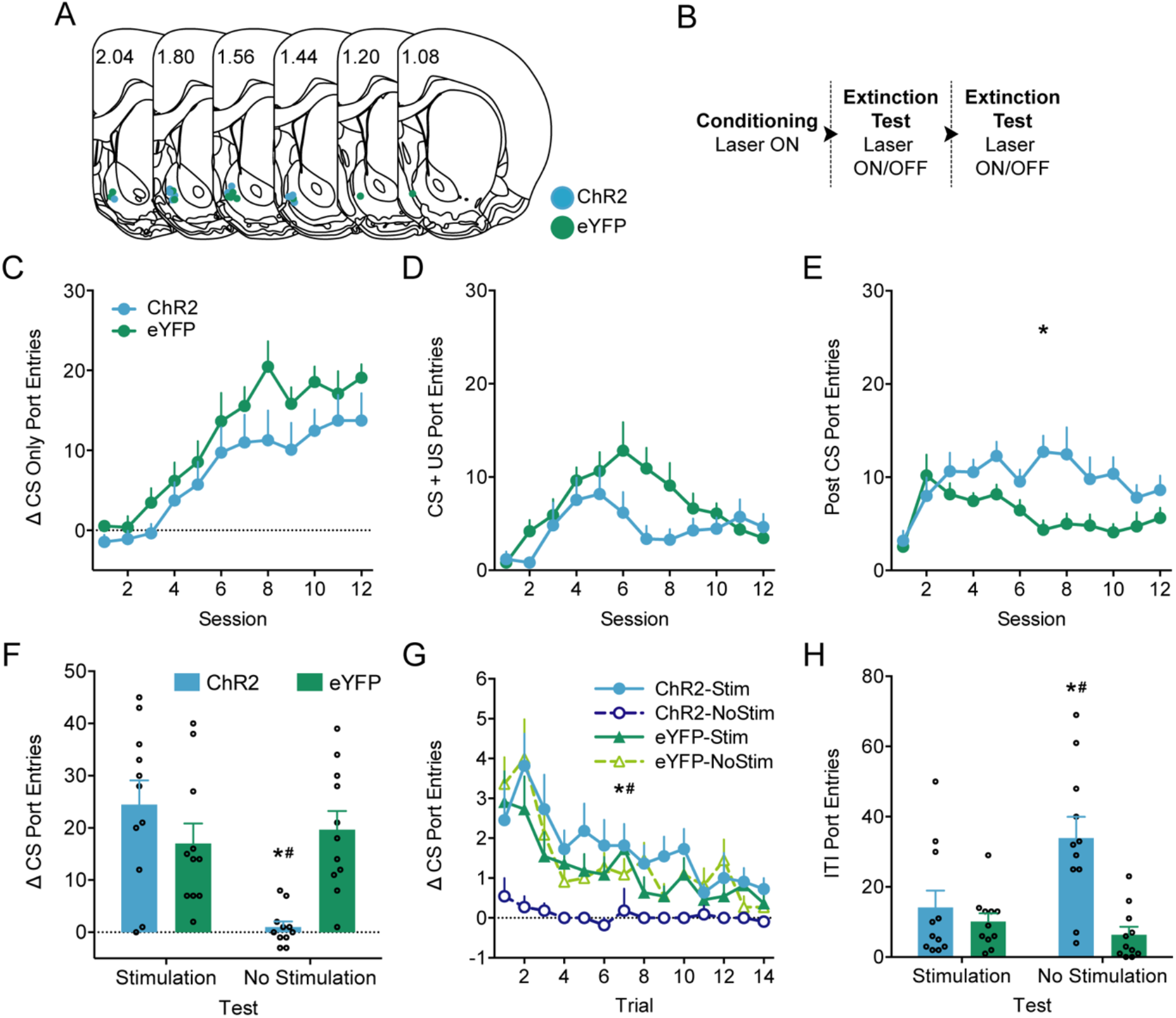
Optogenetic stimulation of the IL-to-NAcS neural circuit altered Pavlovian conditioning. **(A)** Optical fiber placements in the NAcS of rats expressing either ChR2 (blue) or eYFP alone (green) included in the final data analysis. Numbers are locations of sections relative to bregma. **(B)** Design of behavioural procedures. **(C)** Δ CS only port entries (4 sec) across Pavlovian conditioning sessions with IL-to-NAcS stimulation during CS trials. **(D)** Port entries during the 6 sec overlapping CS and US interval across Pavlovian conditioning sessions. (**E**) Post CS port entries across Pavlovian conditioning sessions. * p < 0.05 main effect of virus group. (**F**) Δ CS port entries during the expression test under extinction conditions in the presence or absence of IL-to-NAcS optogenetic stimulation during the CS. (**G**) Δ CS port entries across trials in the expression test under extinction conditions in the presence or absence of optogenetic stimulation during the CS. (**H**) ITI port entries during the expression test under extinction conditions in the presence or absence of optogenetic stimulation during CS trials. (**F-H**) * p < 0.05 ChR2-No Stim vs. eYFP-No Stim. # p < 0.05 ChR2-No Stim vs. ChR2-Stim. All data are mean ± SEM.

During conditioning, the US was initiated 4 seconds after CS onset. Therefore, we analyzed the effect of IL-to-NAcS stimulation on a CS only interval consisting of the first 4 seconds after CS onset, as well as on a 6 second interval encompassing the CS and US during conditioning. A Δ CS only port entry score was calculated by subtracting port entries made 4 s before CS onset from port entries made during the 4 s CS only interval. Δ CS only port entries increased equivalently across conditioning sessions in both the ChR2 and eYFP group (Figure 5C; Session, F(11,220) = 20.18, p < .001; Virus, F(1,20) = 2.58, p = .124, Session x Virus, F(11,220) = 1.37, p = .240). Port entries made during the combined CS and US interval were not affected by IL-to-NAcS stimulation (Figure 5D; Session, F(11,220) = 5.55, p < .001; Virus, F(1,20) = 3.45, p = .078, Session x Virus, F(11,220) = 1.75, p = .143). The trending main effect of virus is likely due to the reduced number of port entries in sessions 6-8 of Pavlovian conditioning in the ChR2 group. Interestingly, post CS port entries (10 s interval after CS offset) were greater in the ChR2 group relative to the eYFP group during Pavlovian conditioning (Figure 5E; Session, F(11,220) = 3.49, p = .007; Virus, F(1,20) = 16.08, p = .001). Further, IL-to-NAcS stimulation decreased probability and increased latency of CS only port entries, with no effect on duration of CS only port entries or ITI port entries (Supplementary Figure 7).

In sum, IL-to-NAcS stimulation did not prevent acquisition of appetitive Pavlovian conditioned responding but may have affected other aspects of CS responding and increased post CS port entries.

### IL-to-NAcS stimulation was required for the expression of appetitive Pavlovian conditioned responding

Following conditioning rats were tested in counterbalanced order for the expression of appetitive Pavlovian conditioned responding under extinction conditions. At test, removing optogenetic stimulation abolished Δ CS port entries in the ChR2 group but not the eYFP group (Figure 5F; Test, F(1,20) = 7.62, p = .012; Virus, F(1,20) = 2.89, p = .105; Test x Virus, F(1,20) = 11.97, p = .002). The eYFP group displayed an equivalent, high number of Δ CS port entries at test in the presence or absence of stimulation (p = .626). In contrast, the ChR2 group had more Δ CS port entries when IL-to-NAcS stimulation was present during the CS compared to when stimulation was removed (p < .001). In the presence of IL-to-NAcS stimulation, there was no statistically significant difference in Δ CS port entries between the ChR2 and eYFP groups (p = .229). However, the ChR2 group made fewer port entries than the eYFP group at test when stimulation was removed (p < .001). A similar pattern of results to Δ CS port entries was observed in post CS port entries, and probability, duration, and latency of CS port entries (Supplementary Figure 8).

In sum, removing IL-to-NAcS stimulation reduced Δ CS port entries in the ChR2 group without affecting responding in the eYFP group.

Analysis of Δ CS port entries per CS trial revealed that removing IL-to-NAcS stimulation abolished responding to the CS from the first trial and onwards in the ChR2 group (Figure 5G; Test, F(1,20) = 7.62, p = .012; Virus, F(1,20) = 2.89, p = .105); Test x Virus, F(1,20) = 11.97, p = .002). This result recapitulates the differences observed in averaged Δ CS port entries between ChR2 and eYFP groups. Within-session extinction was observed as Δ CS port entries decreased across CS trials (Trial, F(13,260) = 10.07, p < .001). However, there were no statistically significant Trial x Virus (F(13,260) = 1.345, p = .224), Trial x Test (F(13,260) = .89, p = .540) or Trial x Test x Virus (F(13,260) = 1.88, p = .050) interactions. The near significant Trial x Test x Virus interaction is likely the result of a reduction in the number of Δ CS port entries, especially in earlier CS trials, in the ChR2 group relative to the eYFP group when stimulation was removed. In contrast, Δ CS port entries underwent within-session extinction and decreased equivalently across trials in both the eYFP group and the ChR2 group when stimulation was present during CS trials.

At test, removing optogenetic stimulation increased ITI port entries in the ChR2 group but not in the eYFP group (Figure 5H; Test, F(1,20) = 5.20, p = .034; Virus, F(1,20) = 10.84, p = .004; Test x Virus, F(1,20) = 11.14, p = .003). The eYFP group showed an equivalent number of ITI port entries in the presence or absence of optogenetic stimulation (p = .464). In contrast, removing IL-to-NAcS stimulation in ChR2 group increased ITI port entries relative to when stimulation was present (p = .001). There was no difference in ITI port entries between the ChR2 and eYFP group when stimulation was present (p = .468). However, when optogenetic stimulation was absent at test, the ChR2 group made more ITI port entries than the eYFP group (p < .001). Therefore, removing IL-to-NAcS stimulation increased port entries made outside the CS in the ChR2 group without affecting responding in the eYFP group.

## DISCUSSION

We report that optogenetically stimulating the IL-to-NAcS neural circuit during CS trials suppressed context-induced renewal of appetitive Pavlovian conditioned responding, without affecting responding in the extinction context or outside CS trials. In a separate experiment, stimulating the IL-to-NAcS circuit suppressed conditioned responding regardless of prior extinction and seemed to impair extinction retrieval. Finally, IL-to-NAcS stimulation altered but did not prevent the acquisition of Pavlovian conditioning and was necessary to maintain the subsequent expression of conditioned responding. These results suggest that stimulating the IL-to-NAcS circuit suppresses responding to appetitive Pavlovian cues and this effect does not require prior extinction.

As predicted, in experiment 1, optogenetically stimulating the IL-to-NAcS circuit during CS trials suppressed the renewal of conditioned responding. This effect occurred in the first CS trial at test in Context A and was reflected in all measures of conditioned responding (Δ CS port entries, probability, latency, duration of CS port entries). These results are consistent with the proposed role of the IL in suppressing appetitive Pavlovian responses (Rhodes and Killcross, 2004; 2007; Villaruel et al., 2018) and with evidence that IL inputs to the NAcS are critical for suppressing operant cocaine-seeking (Peters et al., 2008; 2009; LaLumiere et al., 2012; Augur et al., 2016; Cameron et al., 2019; Warren et al., 2019). Our results extend these findings to appetitive Pavlovian responses using sucrose, a natural reinforcer. Therefore, the suppression of Pavlovian responding to appetitive cues and operant cocaine-seeking after extinction may be mediated by a common neural substrate.

Experiment 2 tested whether IL-to-NAcS stimulation suppressed renewal by promoting the expression of an inhibitory extinction memory. Thus, following Pavlovian conditioning rats received either extinction training or no extinction prior to test. Optogenetically stimulating the IL-to-NAcS circuit at test suppressed CS responding regardless of prior extinction. Therefore, the suppression of renewal in experiment 1 following IL-to-NAcS stimulation may have been accomplished through an extinction-independent process. These results are inconsistent with the hypothesis that IL and IL-to-NAcS circuit stimulation suppresses operant cocaine-seeking by promoting extinction retrieval (Peters et al., 2008; Augur et al., 2016; Ewald Müller et al., 2019). This discrepancy could be related to our use of Pavlovian conditioning, whereas studies investigating the IL-to-NAcS circuit in extinction typically use operant cocaine self-administration. Different psychological processes have been proposed in the extinction of Pavlovian versus operant responding, particularly with regards to the role of context in extinction (Trask et al., 2017), which could influence the recruitment of the IL-to-NAcS circuit by extinction.

Alternately, studies in operant cocaine-seeking used chemogenetics and stable-step function opsins to diffusely enhance IL and IL-to-NAcS activity (Augur et al., 2016; Ewald Müller et al., 2018), whereas we used optogenetics to stimulate the IL-to-NAcS circuit at discrete points during behaviour. Similar application of optogenetics to stimulate the IL and IL-to-NAcS circuit suppressed operant responding for food and cocaine without prior extinction (Do Monte et al., 2015; Cameron et al., 2019). Optogenetic stimulation of IL inputs in the NAcS may disrupt time-locked inhibitory activity in the NAcS that permits consummatory behaviours (Nicola et al., 2004; Taha and Fields, 2006; Krause et al., 2010; Reed et al., 2018), which is a component of the CS-elicited port entry response in our task.

The IL mediates inhibitory associations established through latent inhibition, wherein exposure to a non-reinforced CS delays the subsequent acquisition of Pavlovian conditioning (Lingawi et al., 2016). In our task, exposure to a non-reinforced CS, and therefore the establishment of inhibitory associations through latent inhibition, may have occurred early in conditioning as rats were first learning to enter the fluid port during the CS to obtain sucrose. Pharmacologically stimulating the IL strengthens inhibitory associations acquired through latent inhibition (Lingawi et al., 2016). Therefore, optogenetically stimulating the IL-to-NAcS circuit could potentially suppress responding regardless of prior extinction by facilitating the expression of inhibitory associations established early in Pavlovian conditioning through latent inhibition.

Different neural ensembles involved in promoting and suppressing operant responding have been identified in the IL and in IL projections to the NAcS (Warren et al., 2016; Warren et al., 2019). Further, pharmacologically inactivating the IL can reduce operant food- and heroin-seeking (Bossert et al., 2012; Eddy et al., 2016), suggesting that the IL is also involved in promoting responding after extinction. Therefore, our global stimulation of the IL-to-NAcS circuit may have disrupted the activity of neural ensembles involved in promoting responding, thereby suppressing appetitive Pavlovian conditioned responding regardless of extinction.

In experiment 2, optogenetically stimulating the IL-to-NAcS circuit during extinction did not facilitate extinction retrieval 24 hr later, and in fact, seemed to increase CS responding during the retrieval test in rats with previous extinction training. Therefore, IL-to-NAcS stimulation during extinction seemed to weaken the previously established inhibitory extinction memory. This finding contrasts with studies in aversive conditioning that report facilitated extinction retrieval and strengthening of inhibitory memory after enhancing IL activity during extinction (Milad and Quirk, 2002; Milad, 2004; Do Monte et al., 2015; Lingawi et al., 2016; 2018). These divergent findings could be due to differences in the affective properties of extinction in aversive and appetitive procedures (Amsel, 1958; Gerber et al., 2014).

Optogenetically stimulating the IL-to-NAcS circuit in experiment 3 did not prevent the acquisition or expression of appetitive Pavlovian conditioning, indicating that this manipulation does not result in general motor inhibition. However, stimulation of this neural circuit affected probability and latency measures of conditioned responding and increased post CS port entries, suggesting that conditioning was altered. Optogenetic stimulation of various glutamatergic inputs into the NAcS including the IL supports self-stimulation (Britt et al., 2012; Cameron et al., 2019), suggesting that stimulating this circuit can generate a perceptible stimulus. Therefore, IL-to-NAcS stimulation in experiment 3 may have generated a predictive stimulus that functioned in compound with the white noise CS to signal sucrose delivery. Interestingly, IL-to-NAcS stimulation was required for the subsequent expression of responding to the white noise CS. Removing stimulation decreased CS responding and increased port entries outside of the CS. These results suggest that IL-to-NAcS stimulation acted as a predictive cue for sucrose and altered Pavlovian conditioning.

Although ChR2 expression was predominantly in the IL, we observed some spread in the medial orbitofrontal cortex (OFC) and lateral parts of the prelimbic cortex (PL) along the forceps minor of the corpus callosum. Our results could thus be due to optogenetic stimulation of inputs in the NAcS from regions neighbouring the IL. However, the PL is thought to promote cocaine-seeking (McFarland and Kalivas, 2001; Capriles et al., 2003; McLaughlin and See, 2003) and optogenetic stimulation of the PL-to-NAc circuit elevates Pavlovian conditioned responding (Otis et al., 2017). Furthermore, pharmacologically inactivating the OFC disrupts over-expectation but not extinction in aversive conditioning (Lay et al., 2020). Finally, our findings in experiments 1 and 2 are consistent with numerous studies on the role of the IL and the IL-to-NAcS circuit in extinction using different procedures (Quirk et al., 2000; Milad and Quirk, 2002; Peters et al., 2008; LaLaumiere et al., 2012; Augur et al., 2016; Villaruel et al., 2018), suggesting that we successfully targeted the IL-to-NAcS circuit.

We found that IL projections to the NAcS and BLA were largely made up of anatomically distinct neural subpopulations (Bloodgood et al., 2018). Unilateral optogenetic stimulation of the IL-to-NAcS circuit induced bilateral Fos activation in the IL and the NAcS, which may be due to bilateral projections from the IL to the NAcS and contralateral connections between the IL (Hurley et al., 1991; Vertes, 2004). However, Fos activation was greater in the hemisphere transfected with ChR2 and implanted with the optical fiber, suggesting that the IL-to-NAcS circuit is predominantly ipsilateral.

The present research was conducted only in male rats and is unable to observe sex differences. There are sex differences in the role of the IL in renewal of appetitive Pavlovian conditioned responding (Anderson and Petrovich, 2018a; 2018b). Sex differences in behavioural strategies could lead to differential effects of neural manipulations (Radke et al., 2021; Shanksy and Murphy, 2021). We observe similar levels of renewal (Brown and Chaudhri, 2021 Conference Abstract) and reinstatement (LeCocq et al., 2021 Conference Abstract) of appetitive Pavlovian conditioned responding in male and female rats. Our ongoing research on corticothalamic regulation of extinction is designed to investigate potential sex differences (Brown and Chaudhri, unpublished).

In conclusion, optogenetically stimulating the IL-to-NAcS circuit suppressed the renewal of appetitive Pavlovian conditioned responding and may have done so in an extinction-independent manner. Additionally, IL-to-NAcS stimulation did not facilitate extinction retrieval, which contrasts with findings in aversive conditioning. Importantly, stimulation of this circuit did not suppress motor function, and altered Pavlovian conditioning by potentially acting as a stimulus that became associated with other external stimuli. The present results advance our understanding of the complex processes by which the IL-to-NAcS circuit controls appetitive Pavlovian conditioned responding. Further work is needed in both sexes to determine the mechanisms by which this corticostriatal circuit mediates adaptive behaviour to meet environmental demands. Understanding the corticostriatal mechanisms involved in inhibiting learned responses can aid in the treatment of disorders such as substance abuse and post-traumatic stress.

## FUNDING

The Natural Sciences and Engineering Research Council of Canada (NSERC; 387 224-2010, N.C.) funded this research. N.C. is the recipient of a Chercheur-Boursier Junior 2 award from Fonds de recherche du Québec—Santé (FRQS). F.V. was supported by graduate fellowships from Concordia University and NSERC.

## ACKNOWLEDGEMENTS

N.C. and F.R.V. designed all experiments. F.R.V. conducted and analyzed the data for all experiments and M.M. helped conduct and analyze data for experiment 2. F.R.V. prepared the article with input from N.C. and M.M. All authors reviewed the manuscript and provided comments. We thank Steve Cabilio for support with Med PC programming and data extraction, David Munro for technical support, and Dr. Chris Law of the Centre for Microscopy and Cellular Imaging for writing the FIJI macro and programs for Fos and tracer quantification and technical support in microscopy. Conflicts of interest: The authors declare no conflicts of interest.

## SUPPLEMENTARY MATERIAL

**Supplementary Figure 1.**
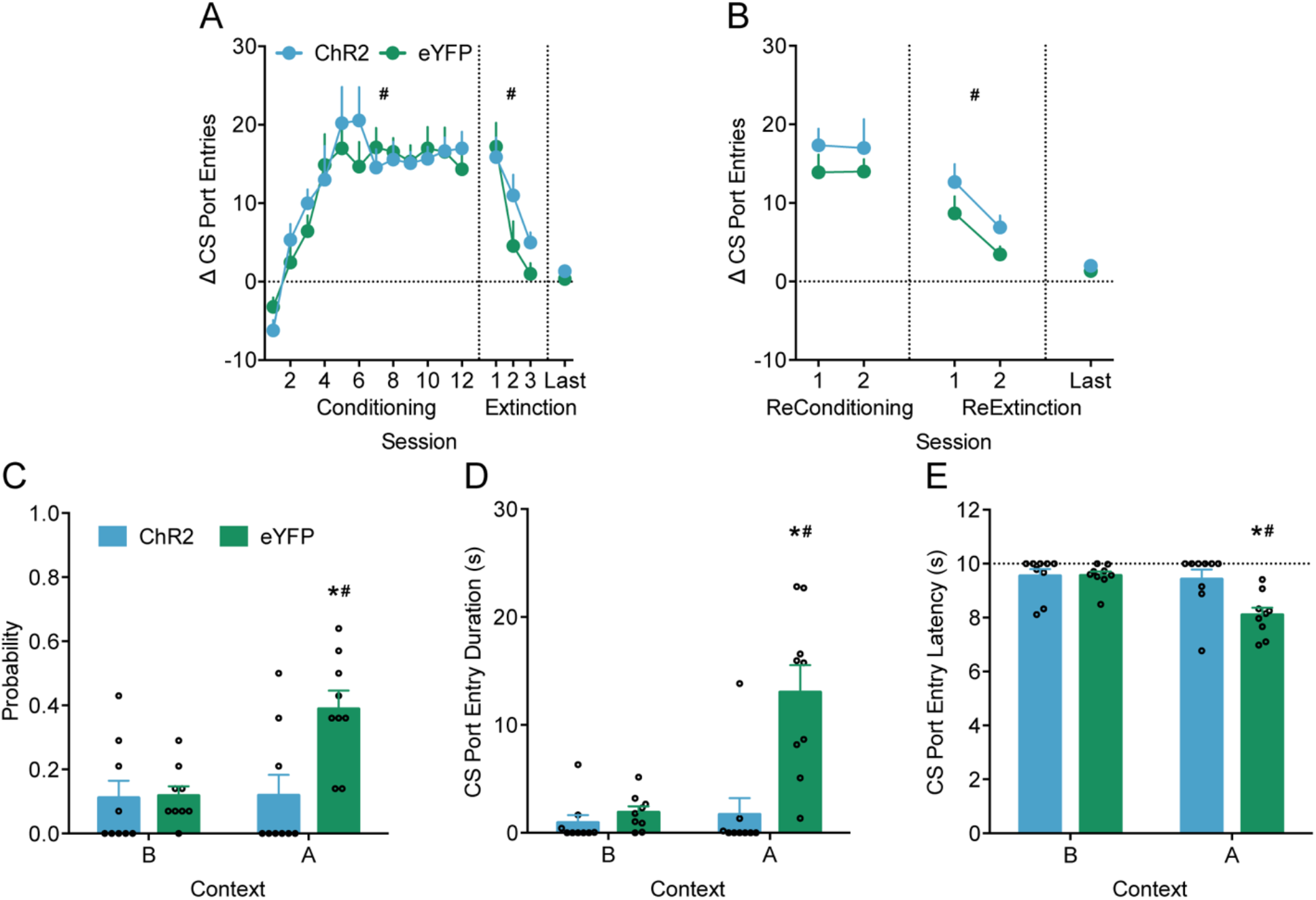
Experiment 1 data for acquisition, extinction, and additional dependent variables at tests. **(A)** Both ChR2 and eYFP groups similarly acquired Pavlovian conditioning in Context A as measured by Δ CS port entries (Session, F(11,176) = 14.59, p < .001; Virus, F(1,16) = .172, p = .684; Session x Virus, F(11,176) = .62, p = .628) and extinguished conditioned responding in the first three sessions of extinction in Context B (Session, F(2,32) = 16.84, p < .001; Virus, F(1,16) = 2.08, p = .168; Session x Virus, F(2,32) = 1.41, p = .259). Virus groups did not differ in the last session of extinction prior to test (Virus, F(1,16) = .51, p = .487). **(B)** Both ChR2 and eYFP groups similarly re-acquired (Session, F(1,16) = .004, p = .949; Virus, F(1,16) = 1.06, p = .319; Session x Virus, F(1,16) = .02, p = .898) and re-extinguished (Session, F(1,16) = 13.67, p = .002; Virus, F(1,16) = 3.16, p = .094; Session x Virus, F(1,16) = .04, p = .854) Pavlovian conditioned responding following the first round of testing. Groups did not differ in Δ CS port entries in the last session of extinction prior to the second round of tests (Virus, F(1,16) = .53, p = .476). (**A-B**) # p < 0.05 main effect of session. (**C**) At tests, the eYFP group displayed renewal with higher probability of CS port entries at test in the conditioning context (Context A) relative to test in the extinction context (Context B). Stimulation of the IL-to-NAcS neural circuit in the ChR2 group attenuated the probability of CS port entries in Context A relative to the eYFP group (Virus F(1,16) = 4.09, p = .060; Context F(1,16) = 21.647, p < .001; Virus x Context F(1,16) = 19.34, p < .001). (**D**) Total duration of port entries initiated during the CS was greater for the eYFP group at test in Context A relative to Context B, indicating renewal of conditioned responding. IL-to-NAcS stimulation attenuated the duration of CS port entries in the ChR2 group relative to the eYFP group in Context A (Virus F(1,16) = 12.57, p = .003, Context F(1,16) = 20.12, p < .001; Virus x Context F(1,16) = 15.26, p = .001). **(E)** Average latency to make a CS port entry was lower at test in Context A relative to Context B in the eYFP group, indicating renewal. Stimulation of the IL-to-NAcS in the ChR2 group increased the latency to make a CS port entry relative to the eYFP group in Context A (Virus F(1,16) = 3.75, p = .071; Context F(1,16) = 19.73, p <.001; Virus x Context F(1,16) = 14.16, p = .002). Dashed line indicates duration of the CS and maximum latency. (**C-E**) # p < 0.05 Context A vs. Context B in the eYFP group. * p < 0.05 ChR2 vs. eYFP in Context A. All data are mean ± SEM.

**Supplementary Figure 2.**
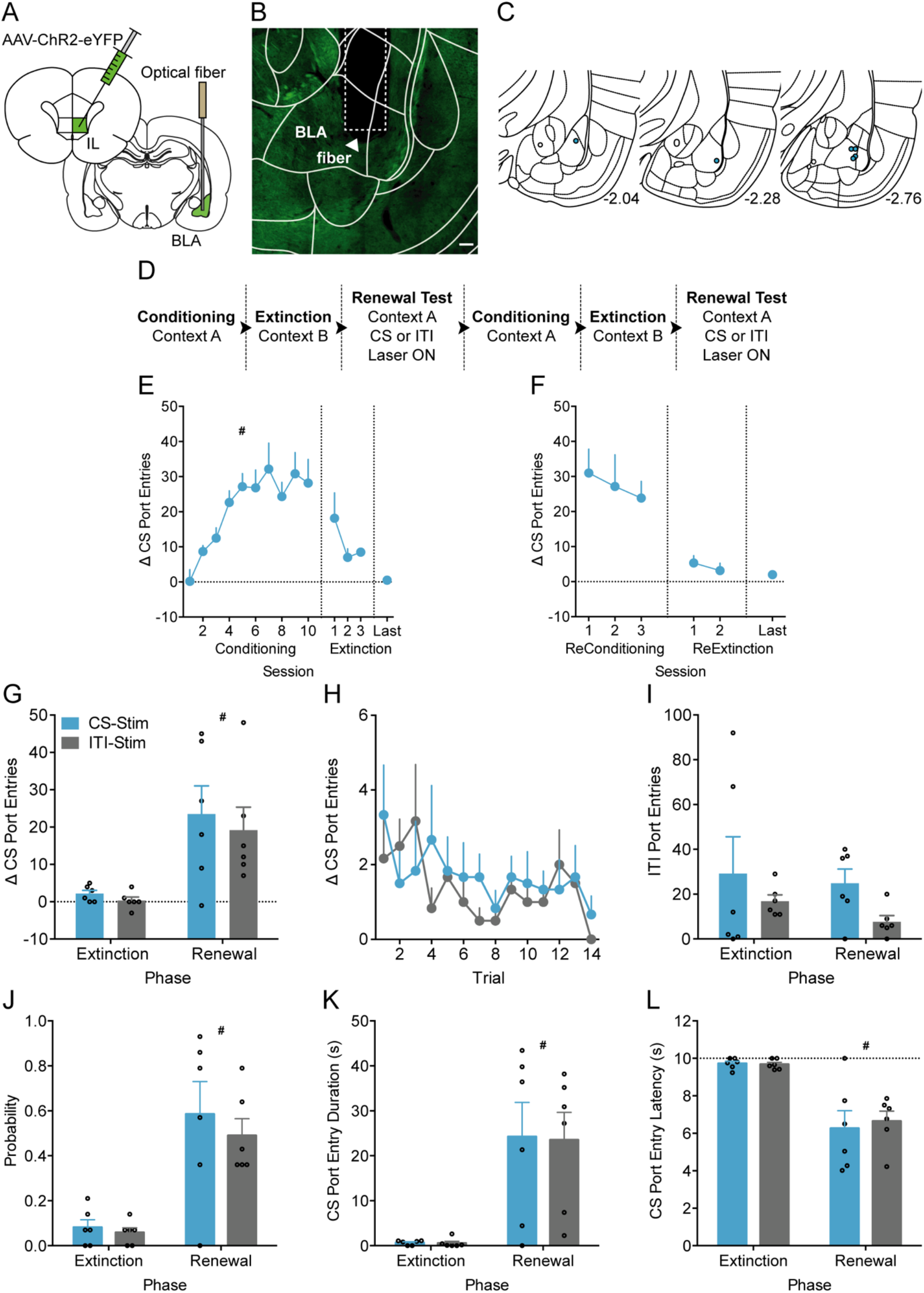
Optogenetic stimulation of the IL-to-Basolateral Amygdala (BLA) neural circuit did not affect context-induced renewal of appetitive Pavlovian conditioned responding. **(A)** Method schematic for microinjection of ChR2 in the IL and an optical fiber implanted in the BLA (AP −2.5 mm, ML −5.0 mm, relative to bregma, DV −8.5 mm relative to skull surface) to target the IL-to-BLA neural circuit. **(B)** Representative image depicting ChR2 expression of IL terminals in the BLA and optical fiber placement. Scale bar 100 μm. **(C)** Optical fiber placements in the BLA for ChR2 (blue) expressing rats included in the final data analysis (n = 6). Numbers are locations of sections relative to bregma. **(D)** Design of behavioural procedures. **(E)** Δ CS port entries increased during conditioning in Context A (Session, F(9,45) = 9.08, p = .002), decreased during extinction in Context B (Session, F(2,10) = 1.58, p = .254) and remained low in the last extinction session prior to the first renewal test. # p < 0.05 main effect of session. **(F)** Δ CS port entries remained high across three reconditioning sessions (Session, F(2,10) = 1.14, p = .359), was lower during the reextinction sessions (Session, F(1,5) = .54, p = .494) and was maintained at low levels in the last extinction session prior to the second renewal test. **(G)** Rats exhibited renewal with greater Δ CS port entries at test relative to the last extinction session regardless of whether optogenetic stimulation of the IL-to-BLA circuit occurred during the CS or in the middle of the intertrial interval (ITI) (Phase, F(1,5) = 10.94, p = .021; Stimulation, F(1,5) = .81, p = .409; Phase x Stimulation, F(1,5) = .114, p = .749). **(H)** Δ CS port entries across trials during the renewal tests were similar regardless of whether IL-to-BLA circuit stimulation occurred during the CS or the ITI (Stimulation, F(1,5) = .38 p = .563; Trial, F(13,65) = 1.49, p = .265; Stimulation x Trial, F(13,65) = .68, p = .593). **(I)** Port entries during the ITI were similar across extinction and renewal regardless of the time of IL-to-BLA stimulation (Phase, F(1,5) = 1.02, p = .359; Stimulation, F(1,5) = 2.94, p = .147; Phase x Stimulation, F(1,5) = .110, p = .754). **(J)** Probability of CS port entries was greater during the renewal test compared to the last extinction session regardless of whether IL-to-BLA stimulation occurred during the CS or the ITI at test (Phase, F(1,5) = 29.48, p = .003; Stimulation F(1,5) = 0.44, p = .538; Phase x Stimulation F(1,5) = 0.27, p = .627). **(K)** Total duration of CS port entries was greater at the renewal test relative to the last extinction session regardless of whether optogenetic stimulation occurred during the CS or ITI (Phase, F(1,5) = 46.72, p = .001; Stimulation F(1,5) = 0.01, p = .948; Phase x Stimulation F(1,5) = 0.003, p = .962). **(L)** Average latency of CS port entries was lower at test compared to the last session of extinction and was not affected by IL-to-BLA stimulation either during the CS or the ITI (Phase, F(1,5) = 30.35, p = .003; Stimulation F(1,5) = 0.08, p = .786; Phase x Stimulation F(1,5) = 0.25, p = .642). Dashed line indicates duration of the CS and maximum latency. **(G, J-L)** # p < 0.05 main effect of phase. All data are mean ± SEM.

**Supplementary Figure 3.**
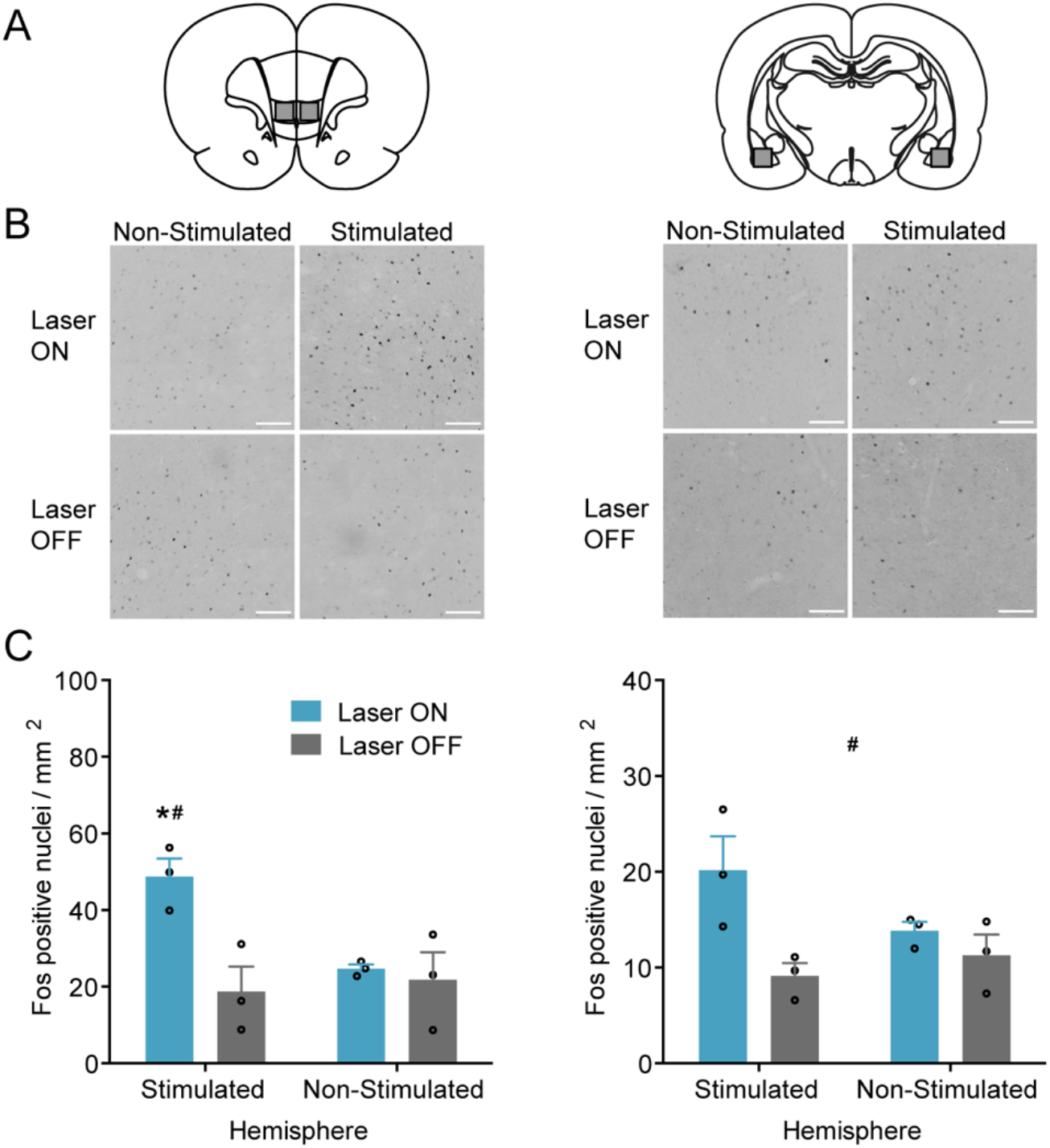
Quantification of Fos positive nuclei density following optogenetic stimulation of the IL-to-BLA neural circuit. (**A**) Schematic depicting areas in images of the IL (left panel) and the BLA (right panel) quantified for Fos positive nuclei. (**B**) Representative images of Fos positive nuclei in the IL (left panel) and the BLA (right panel) in the stimulated and non-stimulated hemisphere of ChR2 expressing rats that received laser stimulation (Laser ON) or no laser stimulation (Laser OFF). Scale bars 100 μm. (**C**) Density of Fos positive nuclei in the IL (left graph) (Hemisphere, F(1,4) = 20.38, p = .011; Stimulation, F(1,4) = 5.00, p = .089; Hemisphere x Stimulation, F(1,4) = 34.09, p = .004). Density of Fos positive nuclei in the IL was greater in the stimulated hemisphere containing the optical fiber in rats that received laser stimulation (Laser ON Stimulated) in comparison to the same hemisphere of rats that did not receive laser stimulation (Laser OFF Stimulated). # p < 0.05 Laser ON vs. OFF in the stimulated hemisphere. Density of Fos positive nuclei was also greater in the stimulated hemisphere (Laser ON Stimulated) than the non-stimulated hemisphere (Laser ON Non-Stimulated) in the same rats. * p < 0.05 stimulated vs. non-stimulated hemisphere in the laser ON condition. Density of Fos positive nuclei in the BLA (right graph) was greater in rats that received laser stimulation compared to rats that did not receive laser stimulation (Stimulation, F(1,4) = 8.73, p = .042; Hemisphere, F(1,4) = .96, p = .384; Hemisphere x Stimulation F(1,4) = 3.88, p = .120). # p < 0.05 main effect of laser stimulation. All data are mean ± SEM.

**Supplementary Figure 4.**
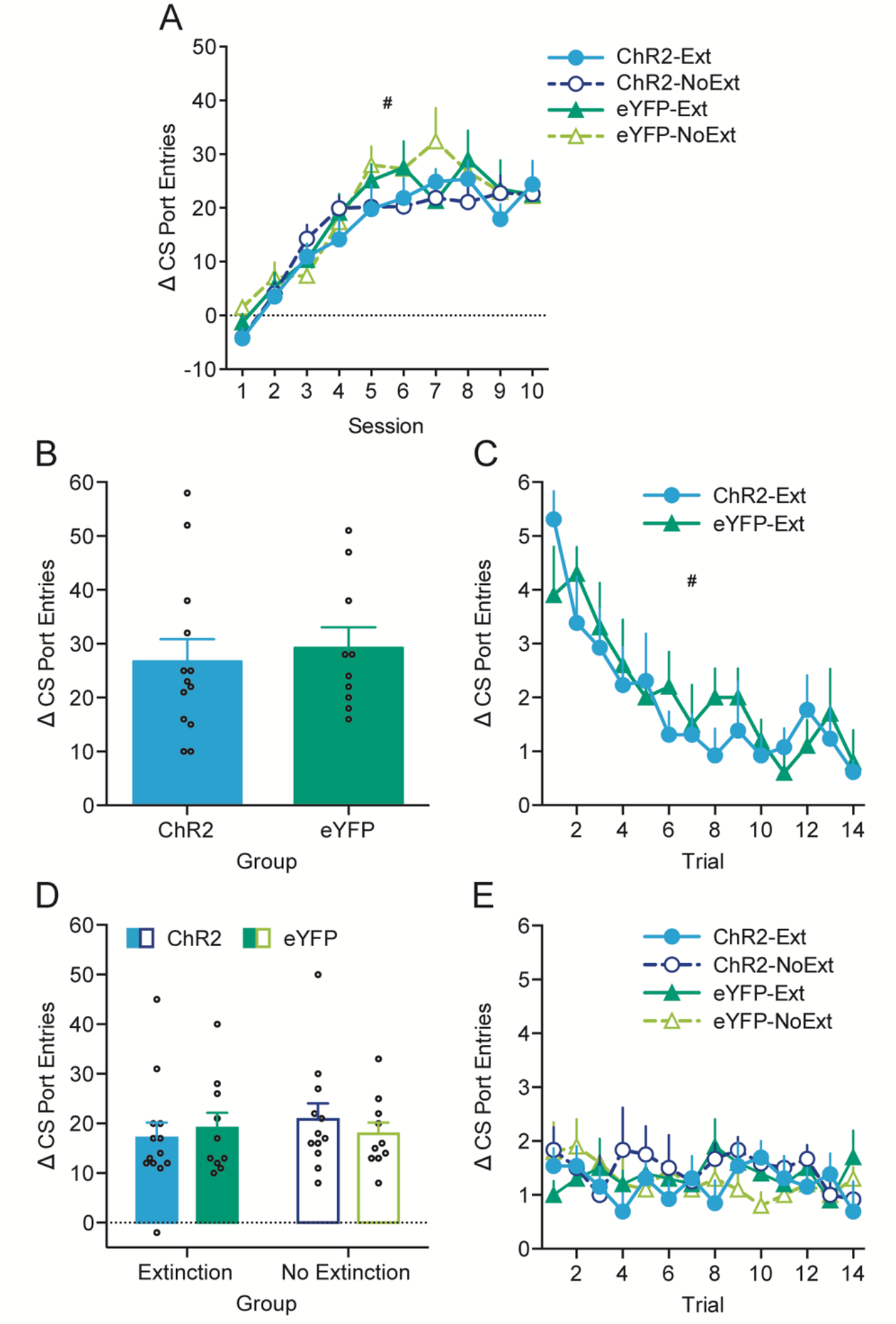
Experiment 2 Δ CS port entries across Pavlovian conditioning, extinction, and reconditioning sessions. **(A)** All groups displayed an equivalent increase in Δ CS port entries across conditioning sessions (Session, F(4,161) = 41.52, p = .001; Virus, F(1,41) = 1.90, p = .176; Group F(1,41) = 0.14, p = .708; Session x Virus F(4,161) = 1.17, p = .326; Session × Group, F(4, 161) = 0.48, p = .747; Virus × Group, F(1, 41) = 0.03, p = .872; Session × Virus × Group, F(3.93, 161.28) = 1.06, p = .376). # p < 0.05 main effect of session. **(B)** Within the Extinction group, overall Δ CS port entries were equivalent between ChR2 and eYFP groups (Virus, F(1,21) = .19, p = .670). **(C)** ChR2 and eYFP Extinction groups had similar within-session reduction in Δ CS port entries across trials during the extinction session (Trial, F(13, 273) = 7.30, p < .001; Virus, F(1, 21) = 0.19, p = .670; Trial x Virus, F(13, 273) = 0.66, p = .692). # p < 0.05 main effect of trial. **(D)** Δ CS port entries during the reconditioning session was comparable across all virus and extinction groups (Virus, F(1, 41) = 0.02, p = .881; Group, F(1, 41) = 0.18, p = .675; Virus x Group, F(1, 41) = 0.67, p = .417). **(E)** Δ CS port entries across trials during the reconditioning session was similar across virus and extinction groups (Trial, F(13,533) = 0.67, p = .724; Virus, F(1,41) = 0.02, p = .881; Group, F(1,41) = 0.18, p = .675; Virus x Group, F(1,41) = 0.67, p = .417; Trial x Virus, F(13,533) = 0.96, p = .471; Trial x Group, F(13,533) = 0.57, p = .814; Trial x Virus x Group, F(13,533) = 0.80, p = .610). All data are mean ± SEM.

**Supplementary Figure 5.**
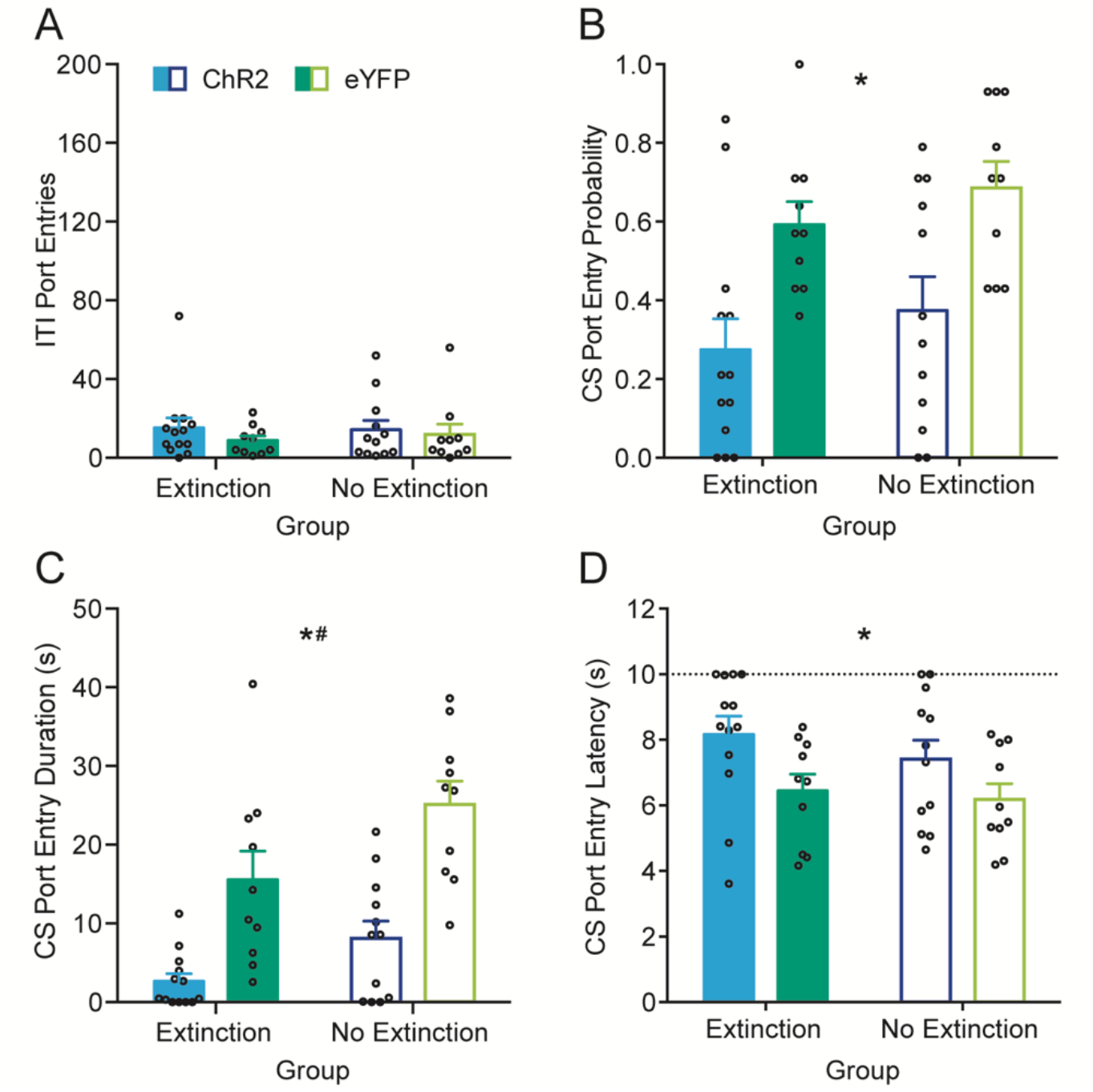
Experiment 2 additional dependent variables during the extinction test (Test 1). Optogenetic stimulation of the IL-to-NAcS circuit during CS trials did not affect port entries during the intertrial interval (ITI) but attenuated CS port entry probability and duration, and increased latency regardless of prior extinction training. **(A)** ITI port entries was equivalent across all groups (Virus, F(1,41) = 0.91, p = .346; Group, F(1,41) = 0.05, p = .829; Virus x Group F(1,41) = 0.18, p = .671). **(B)** IL-to-NAcS stimulation attenuated the probability of CS port entries in the ChR2 groups relative to the eYFP groups regardless of prior extinction training (Virus, F(1,41) = 17.08, p < .001; Group, F(1,41) = 1.61, p = .211; Virus x Group F(1,41) = .002, p = .961). **(C)** IL-to-NAcS stimulation attenuated total duration of CS port entries in the ChR2 groups relative to the eYFP groups. Across virus groups, the Extinction group displayed lower CS port entry durations relative to the No Extinction group. (Virus, F(1,41) = 36.26, p < .001; Group, F(1,41) = 9.15, p = .004; Virus x Group F(1,41) = .69, p = .412). **(D)** Optogenetic stimulation of the IL-to-NAcS circuit increased the average latency to initiate a CS port entry in the ChR2 group relative to the eYFP group regardless of prior extinction (Virus, F(1,41) = 7.19, p = .011; Group, F(1,41) = .86, p = .360; Virus x Group F(1,41) = .21, p = .651). **(B-D)** * p < 0.05 main effect of virus. # p < 0.05 main effect of extinction group. All data are mean ± SEM.

**Supplementary Figure 6.**
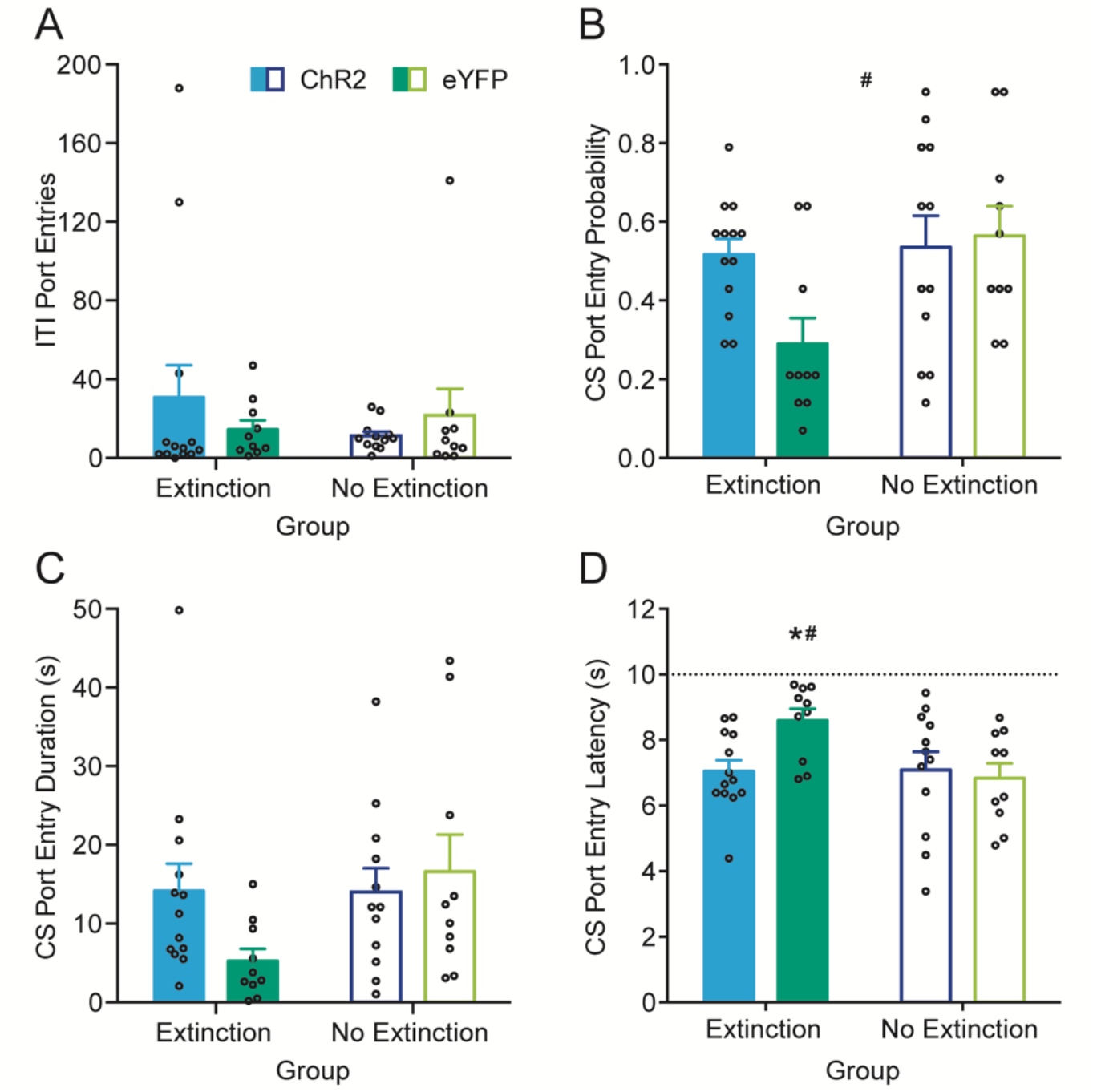
Experiment 2 additional dependent variables during the extinction retrieval test (Test 2). Extinction retrieval seemed to be impaired following optogenetic stimulation of the IL-to-NAcS neural circuit during the extinction test (Test 1) in rats that previously received extinction training. **(A)** ITI port entries were not affected by prior optogenetic stimulation of the IL-to-NAcS circuit and was equivalent across all groups during extinction retrieval (Virus, F(1,41) = 0.06, p = .803; Group, F(1,41) = 0.28, p = .597; Virus x Group F(1,41) = 1.34, p = .254). **(B)** Probability of CS port entries during extinction retrieval was lower in the Extinction groups relative to rats that did not receive prior extinction training (No Extinction) (Virus, F(1,41) = 2.20, p = .145; Group, F(1,41) = 4.89, p = .033; Virus x Group F(1,41) = 3.68, p = .062). # p < 0.05 main effect of extinction group. Visual inspection suggests that this effect may have largely been driven by lower probability of CS port entries in eYFP Extinction group compared to the eYFP No Extinction group. The ChR2 Extinction and ChR2 No Extinction groups had similar average probability of CS port entries during extinction retrieval. Within the Extinction group, the ChR2 group displayed higher probability of CS port entries compared to the eYFP group. **(C)** Total duration of CS port entries was similar across all groups during extinction retrieval (Virus, F(1,41) = .86, p = .358; Group, F(1,41) = 2.73, p = .106; Virus x Group F(1,41) = 2.88, p = .097). Visual inspection indicates that the eYFP Extinction group had lower CS port entry durations than the eYFP No Extinction group. The ChR2 Extinction and ChR2 No Extinction groups had similar total duration of CS port entries during extinction retrieval. Within the Extinction group, the ChR2 group showed higher duration of CS port entries relative to the eYFP group. **(D)** IL-to-NAcS stimulation during extinction impaired subsequent extinction retrieval as measured by average latency of initiate a CS port entry in rats that previously received extinction training (Virus, F(1,41) = 2.19, p = .146; Group, F(1,41) = 3.83, p = .057; Virus x Group F(1,41) = 4.22, p = .046). The eYFP Extinction group had greater CS port entry latency than the eYFP No Extinction group. # p < 0.05 Extinction vs. No Extinction in the eYFP group. There was no statistically significant difference in average latency to initiate a CS port entry between the ChR2 and eYFP No Extinction groups and between the ChR2 Extinction and ChR2 No Extinction groups. Within the Extinction group, the ChR2 group showed faster latency to initiate CS port entries relative to the eYFP group, suggesting that prior IL-to-NAcS stimulation during extinction impaired subsequent extinction retrieval. * p < 0.05 ChR2 vs. eYFP in the Extinction group. Dashed line indicates duration of the CS and maximum latency. All data are mean ± SEM.

**Supplementary Figure 7.**
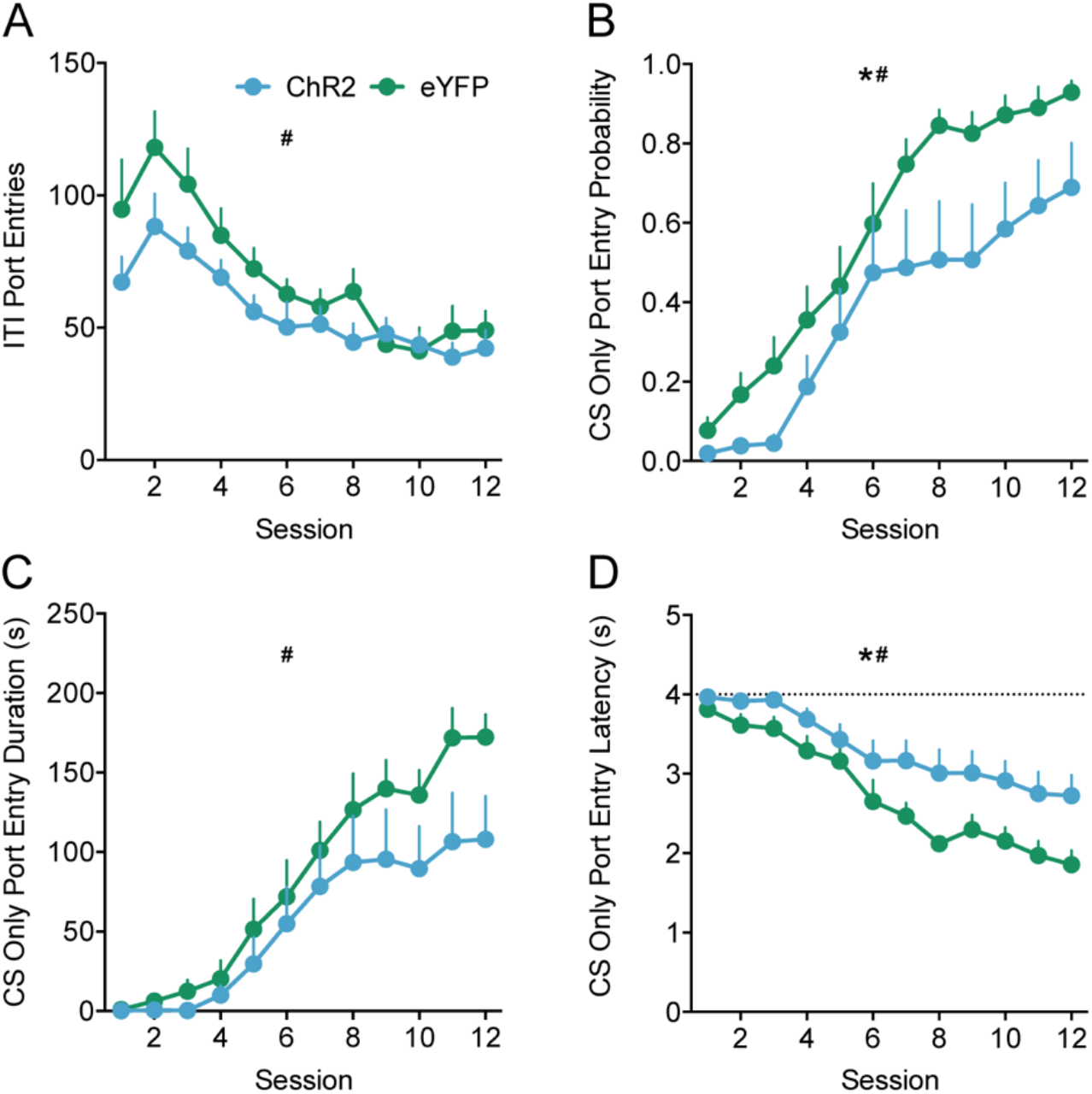
Experiment 3 additional dependent variables during Pavlovian conditioning. Optogenetic stimulation of the IL-to-NAcS altered the acquisition of appetitive Pavlovian conditioning. **(A)** ITI port entries was similar between ChR2 and eYFP groups during Pavlovian conditioning (Session, F(11,220) = 14.75, p < .001; Virus, F(1,20) = 2.81, p = .109; Session x Virus, F(11,220) = 1.07, p = .379). **(B)** Optogenetic stimulation of the IL-to-NAcS circuit during CS trials in conditioning decreased the probability of CS only port entries in the ChR2 group relative to the eYFP group (Session, F(11,220) = 44.51, p < .001; Virus, F(1,20) = 4.49, p = .047; Session x Virus, F(11,220) = 1.14, p = .342). **(C)** Optogenetic stimulation of the IL-to-NAcS circuit during conditioning did not affect the total duration of CS only port entries (Session, F(11,220) = 37.78, p < .001; Virus, F(1,20) = 1.87, p = .187; Session x Virus, F(11,220) = 1.53, p = .229). **(D)** Optogenetic stimulation of the IL-to-NAcS circuit during conditioning increased the latency to initiate CS only port entries in the ChR2 group relative to the eYFP group (Session, F(11,220) = 42.18, p < .001; Virus, F(1,20) = 6.73, p = .017; Session x Virus, F(11,220) = 2.03, p = .104). Dashed line indicates duration of the CS only period and maximum latency. **(A-D)** # p < 0.05 main effect of session. * p < 0.05 main effect of virus. All data are mean ± SEM.

**Supplementary Figure 8.**
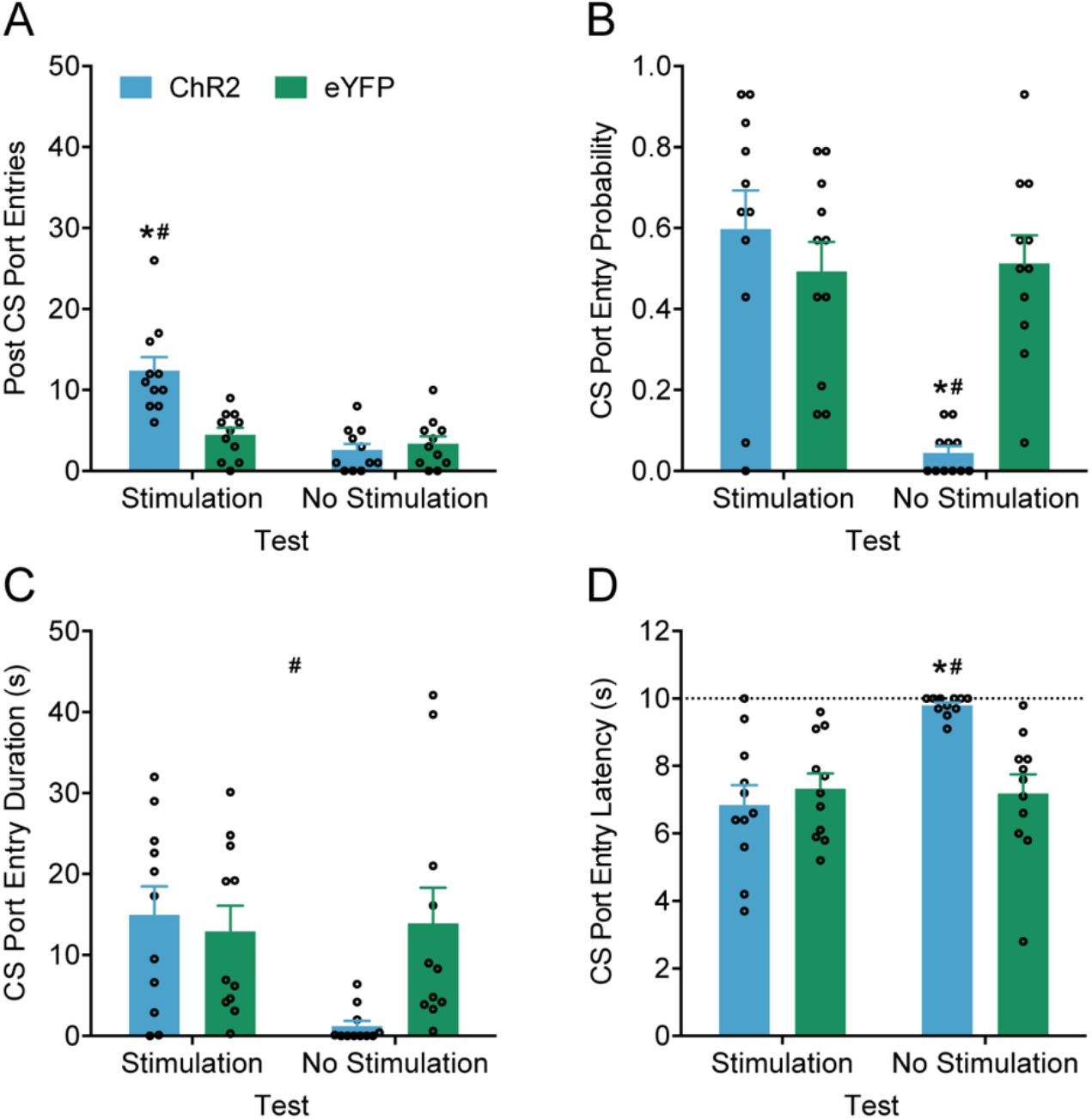
Experiment 3 additional dependent variables during the expression test under extinction conditions in the presence or absence of optogenetic stimulation. Removal of IL-to-NAcS circuit stimulation impaired the expression of appetitive Pavlovian conditioned responding in the ChR2 group but not the eYFP group. **(A)** Post CS port entries during the expression tests remained low in the eYFP group regardless of the presence (Stimulation) or absence of (No Stimulation) optogenetic stimulation. In the ChR2 group, post CS port entries were elevated in the Stimulation expression test and remained low in the No Stimulation test (Test, F(1,20) = 22.32, p < .001; Virus, F(1,20) = 10.35, p = .004; Test x Virus, F(1,20) = 14.28, p = .001). **(B)** Probability of CS port entries remained high in the eYFP group during both the Stimulation and No Stimulation tests. In the ChR2 group, probability of CS port entries was maintained in the Stimulation expression test but was abolished in the No Stimulation test (Test, F(1,20) = 15.08, p = .001; Virus, F(1,20) = 6.42, p = .020; Test x Virus, F(1,20) = 17.43, p < .001). **(C)** Total duration of CS port entries was lower at test in the ChR2 group compared to the eYFP group (Test, F(1,20) = 2.76, p = .113; Virus, F(1,20) = 4.49, p = .047; Test x Virus, F(1,20) = 3.69, p = .069). Visual inspection suggests that this effect may have largely been driven by lower duration of CS port entries in the ChR2 group relative to the eYFP group in the No Stimulation test. # p < 0.05 main effect of virus. **(D)** Average latency of CS port entries remained low in the eYFP group in both the Stimulation and No Stimulation expression tests. In the ChR2 group, average latency of CS port entries was maintained to low levels as the eYFP group in the Stimulation test but increased in the No Stimulation expression test (Test, F(1,20) = 9.94, p = .005, Virus, F(1,20) = 4.72, p = .042; Test x Virus, F(1,20) = 11.95, p = .002). **(A-B**,**D)** * p < 0.05 ChR2-Stim vs eYFP-Stim, # p < 0.05 ChR2-Stim vs ChR2-No Stim. All data are mean ± SEM.

## Notes

### Competing Interest Statement

The authors have declared no competing interest.

